# Menin regulates androgen receptor- and MLL-driven condensation, upsetting regulation of cellular AR-driven transcription

**DOI:** 10.1101/2023.08.04.551977

**Authors:** Junaid Ahmed, Jolyon K Claridge, Attila Meszaros, Peter Tompa

**Author notes:** to whom correspondence should be addressed at, and.

## Abstract

Menin is a protein that is regulated via protein-protein interactions by different binding partners, such as mixed lineage leukemia protein (MLL) and androgen receptor (AR). We observed that menin-AR and menin-MLL interactions are regulated by concentration-dependent dimerization of menin, and its interaction with cancer-related AR. As a result of its oligomerization-dependent interaction with both AR and MLL, menin is recruited into AR-RNA and MLL-RNA condensates formed by liquid-liquid phase separation (LLPS), with different outcomes under AR-overexpression or MLL-overexpression conditions representing different cancer types. At high concentrations promoting menin dimerization, it inhibits MLL-RNA LLPS, while making AR-RNA condensates less dynamic, i.e., more gel-like. Regions of AR show both negative/positive cooperativity in menin binding. AR contains a specific menin-binding region (MBR) in its intrinsically disordered N-terminal domain (NTD), menin binding of which is inhibited by the adjacent DNA-binding domain (DBD), but facilitated by a hinge region located between its DBD and ligand-binding domain (LBD) as well as by N terminus of AR. Interestingly, the hinge region reduces the propensity of full-length AR to undergo LLPS in the presence of RNA, which is facilitated by an alternative hinge region present in the tumor-specific AR isoform, AR-v7. As both menin and MLL are recruited into AR-driven, functional cellular condensates aggravated in the case of AR-v7, we posit that the menin-AR-MLL system represents a fine-tuned condensate module of transcription regulation that is balanced toward the tumor-suppressor activity of menin. Our results suggest that this balance can be upset by prevalent oncogenic events, such as menin upregulation and/or AR-v7 overexpression, in cancer.

## Introduction

Prostate cancer (PC) is the most common cancer in men, against which androgen deprivation therapy (ADT) is the first line of treatment [1]. ADT, however, is often followed by relapse, resulting in a more severe form of cancer termed castration-resistant prostate cancer (CRPC), which is challenging to treat. The primary molecular factor in PC is androgen receptor (AR), a transcription factor that can act as a repressor or activator tuned by coregulators [2]. AR is composed of three structural-functional domains, an intrinsically disordered N-terminal domain (NTD) encompassing trans-activator functionality, a DNA binding domain (DBD), and a ligand binding domain (LBD). The LBD binds activating hormones testosterone and dihydrotestosterone, making AR translocate from the cytoplasm to the nucleus and bind androgen response elements (AREs) in the promoter region of genes, such as prostate-specific antigen (PSA). In CRPC, splice variants of AR, most notably AR-v7, appear, which lack the LBD and thus conveys hormone-independent transcriptional activity [3]. Interestingly, the disordered hinge region of AR connecting DBD and LBD is also replaced in AR-v7 by alternative splicing; this “alt hinge” appears to convey important regulatory input on the flanking domains.

AR also interacts with a range of protein co-regulators, such as menin, a scaffold protein that can act as either activator or repressor of AR function [4]. Upon interaction with AR, menin provides a link to the large scaffold protein mixed lineage leukemia protein (MLL), which is important for transcription, but, when increased in expression, is associated with leukemia and PC [5]. Menin can act as both an oncogene and a tumor suppressor, for example, its inactivation in endocrine glands leads to multiple endocrine neoplasia (MEN) [5,6] caused by reduced antiproliferative gene expression [6]. In leukemia and prostate cancer, it primarily acts as an oncogene [7].

MLL belongs to the histone methyltransferase (HMT) family of enzymes, and it has several isoforms (MLL1, 2, 3 and 4), which catalyze histone H3 Lys4 (K4) mono-, di-, and trimethylation, regulating gene expression [8]. MLL complexes consist of highly conserved core proteins including MLL, WDR5, RBBP5, and ASH2L, which are required for enzymatic activity of complexes. Menin can interact with different types of MLL that derives its importance from the fact that the translocation of MLL gene results in fusion proteins causing acute leukemia via aberrant transcriptional activity. Activity of these chimeric proteins depend on their direct interaction with menin [4,5]. Because of the basic importance of AR-menin-MLL interaction in cell physiology and disease, the interaction between menin and MLL has been characterized to some detail, and small-molecule inhibitors of the interaction are in clinical trials [9–11]. Because of very tight menin-MLL interaction, its targeting is challenging though, and leaves the possibility of specific targeting of the more transient menin-AR interaction open. This interaction is much less characterized, with some limited information coming from *in vitro* pull-down assays, in which AR NTD region G470-T559 was shown to bind menin. Whereas this interaction is important for the transcriptional activity of AR, it has not yet been characterized in detail. It has been shown, however, that in CRPC, the inhibition of the menin-MLL interaction does not necessarily arrest transcription, it rather requires the inhibition of the menin-AR interaction [5].

The complex regulatory interplay between AR, menin, and MLL may entail novel dimensions via recent observations that AR undergoes biomolecular condensation (liquid-liquid phase separation, LLPS) at gene enhancers, of which the DBD is a key driver [12,13]. AR is recruited to super-enhancers and forms condensates [12–14] which may also be implicated in disease. Here we show that menin is recruited to MLL and AR condensates, the menin binding motif (MBM) of MLL phase separates in the presence of RNA, and menin can compete with RNA for MLL binding, inhibiting phase separation. With regards to AR, it also undergoes phase separation, forming condensates into which menin is recruited. As we found that menin can form dimers, while MLL and AR have non-overlapping, different binding sites on menin, the interactions of the two proteins enables complex modes of local topological organization apparent at AR-overexpression or MLL-overexpression conditions. To provide a comprehensive model for the complex regulation of the menin-AR-MLL system, we suggest that targeting MLL complexes may be achieved by the combined inhibition of the interaction between menin-AR and menin-MLL. This model also showcases a new opportunity to target proteins at the condensate level, opening a novel avenue for the targeting of thus-far undruggable transcription factors [15].

## Results

### 1) Interdomain allostery in AR binding to menin

AR is composed of three domains: the N-terminal domain (NTD) followed by a DNA-binding domain (DBD) and a ligand binding domain (LBD). By predicting structural disorder by the IUPred algorithm [16,17], the NTD is intrinsically disordered, followed by two globular domains, DBD and LBD (Fig. 1A). This structural assessment also agrees with AlphaFold prediction (AF-p10275-f1, cf. Fig. 1B), in which NTD prediction is of low confidence that often correlates with structural disorder, also supported by direct biophysical observations [18–21].

**Figure 1.**
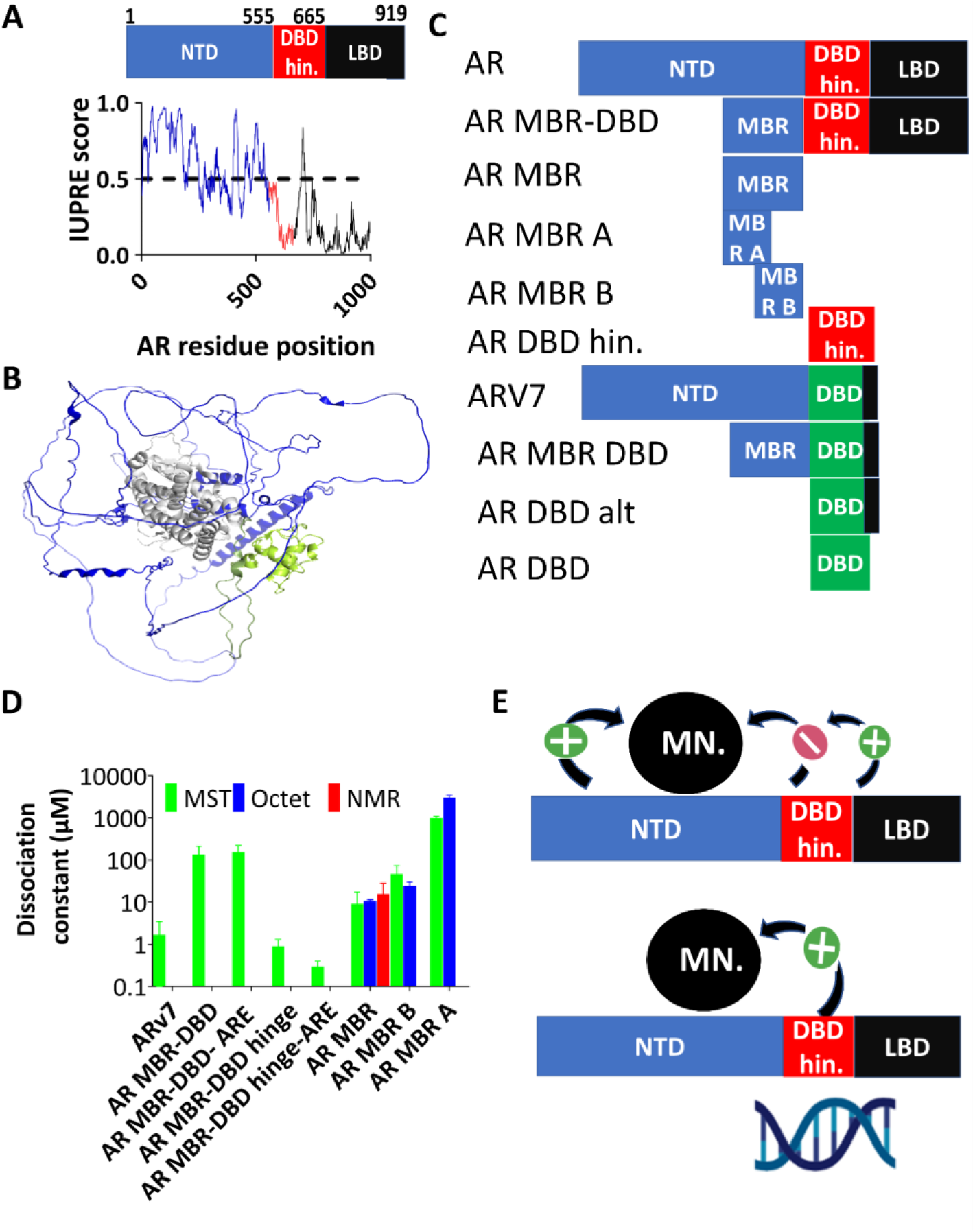
Interaction of menin and androgen receptor. A) Domain architecture of AR along with its disorder tendency predicted by IUPred. Domains shown are the N-terminal domain (NTD), DNA-binding domain (DBD), followed by a short, disordered hinge region (residues 624-655), and the ligand-binding domain (LBD) B) AR structure predicted by AlphaFold (AF-p10275-f1) C) Different constructs of AR, including splice variant AR-v7, AR menin-binding region-DNA binding domain (MBR-DBD), AR menin-binding region (AR MBR), MBR-A and MBR-B. D) The interaction of these constructs with menin with different constructs of AR (for MBR-DBD, with and without ARE), Kd measured by BLI, MST, and NMR. E) Model for allosteric regulation of menin-AR interaction.

It has been reported by immunoprecipitation that AR can interact with menin via a region within its NTD [5], which we here denote as the menin-binding region (MBR, G470-T559) of AR. The strength of direct interaction, as well as the possible allosteric influence of other AR domains on the menin-AR interaction, have not been explored yet. The interaction was only established for full-length (FL) AR; its disease-linked variants, such as AR-v7, have also not been studied in detail. To dissect details of AR-menin interaction, we generated AR constructs of different domain combinations (Fig. 1C).

As AR-v7 is the main driver of CRPC, expressed in 94% of cases and overexpression of menin results in cancer progression, whereas its knock-out has a suppressor effect [22,23], we first addressed the menin ΔIDR1 - AR-v7 interaction by microscale thermophoresis (MST), which shows a Kd = 1.7 ± 1.8 μM (Fig. 1D, Suppl. Fig. S1A). The reason for using this construct was the instability of menin. As the menin-AR interaction was ascribed to the MBR of AR, we next characterized the interaction of menin with the construct AR-MBR-DBD. We found that it binds menin ΔIDR1 much weaker than AR-v7 in the absence of ARE DNA (Kd = 134 ± 77 μM, Fig. 1D, Suppl.Fig. S1B), and slightly weaker in the presence of ARE DNA (Kd = 155 ± 66 μM, Suppl. Fig. S1C). Next, AR-MBD-DBD-LBD interaction was measured with menin to obtain Kd = 0.9 ± 0.4 μM, if AR-MBD-DBD-LBD is complexed with ARE the interaction with menin becomes even tighter (Kd = 0.3 ± 0.1 μM), probably because ARE triggers AR dimerisation, providing extra avidity for interaction with menin (cf. Fig. 1D). The difference in the interaction of menin with AR-v7 and MBR (cf. Fig. 1D) can be explained by two scenarios: either DBD inhibits a direct interaction of menin, or MBR has a weaker affinity than AR-v7, or both. To address the possible allosteric coupling between MBR and DBD, we also measured the interaction of menin with MBR alone. Interestingly, this Kd is much tighter than that of MBR-DBD by either MST (Kd = 6.3 ± 2 μM), Bio-layer interferometry (BLI) (Kd = 10.5 ± 0.9 μM), or NMR (Kd = 15.7 ± 12.5 μM, Fig. 1D, Suppl. Fig. S1 D, E, F).

These differences suggest that DBD inhibits the interaction of menin with MBR, and the NTD has either a direct interaction with menin or allosterically affects menin-MBR interaction, making AR-v7 interaction tighter than with MBR alone.

To conclude we propose a model for allosteric regulation of direct interaction of menin with AR. As menin interacts directly with AR MBR, this interaction is regulated by DBD, Hinge and NTD. DBD inhibits this interaction, whereas Hinge and NTD facilitate this interaction (Fig. 1E).

### 2) Exact binding region of menin within AR

To understand the strong allosteric coupling between MBR and DBD of AR, we next aimed to determine the exact binding region of menin within AR MBR. For this purpose, two separate constructs representing the first and second half of MBR suggested [5] were generated (MBR-A: G473-T518, MBR-B: V509-T559, cf. Fig. 1D), and their interaction with menin was measured. MST and BLI showed a Kd = 24 ± 6 μM and 47 ± 26 μM (MBR-B), and Kd = 1.0 ± 0.1 mM and 3.0 ± 0.4 mM (MBR-A, Fig. 1D, Suppl. Fig S1G, H, I), respectively. These values suggest that the primary menin-binding region of AR falls within MBR-B. To assess if this region binds to the globular domain of menin (M1-V460), to its C-terminal intrinsically disordered region (IDR, R460-P519 (IDR1) and A586-L615 (IDR2)), or both, we also generated a menin ΔIDR1 from which the C-terminal IDR was removed (R460-P519) and a ΔIDR1, IDR2 in which both R460-P519 and Q586-L615 were deleted [24] (keeping the helix linking the two IDRs, IDR1 and IDR2), as it was important for solubility of the construct (cf. Fig. 3B)). The three constructs shows Kds with AR MBR, menin (Kd= 1.7 ± 0.5 μM by MST), menin ΔIDR1 (Kd =6.3 ± 2.0 μM by MST), and menin ΔIDR1, IDR2 (Kd = 2 ± 0.9 μM(by MST) (cf. Suppl. Fig. S1A, J,K) suggesting that the binding region within menin falls in the central folded domain.

To further dissect the menin-AR interaction, we next recorded the ^15^N BEST-HSQC spectrum of ^15^N-labeled AR MBR, which showed a low dispersion in the proton dimension, confirming the intrinsically disordered nature of this region [25]. Small-angle X-ray scattering (SAXS) was used to determine the radius of gyration of MBR (Rg = 2.4 nm), having a Kratky plot with a dip which indicates a disordered protein with some structured /compacted region (s) (Suppl. Fig. S2A) [26]. As the molar mass corresponding to a globular protein of similar Rg is 11.9 kDa, this suggests an extended radius for a 9.4 kDa protein, also arguing for the structural disorder of MBR (Fig. 2D, Suppl. Fig. S2A).

**Figure 2.**
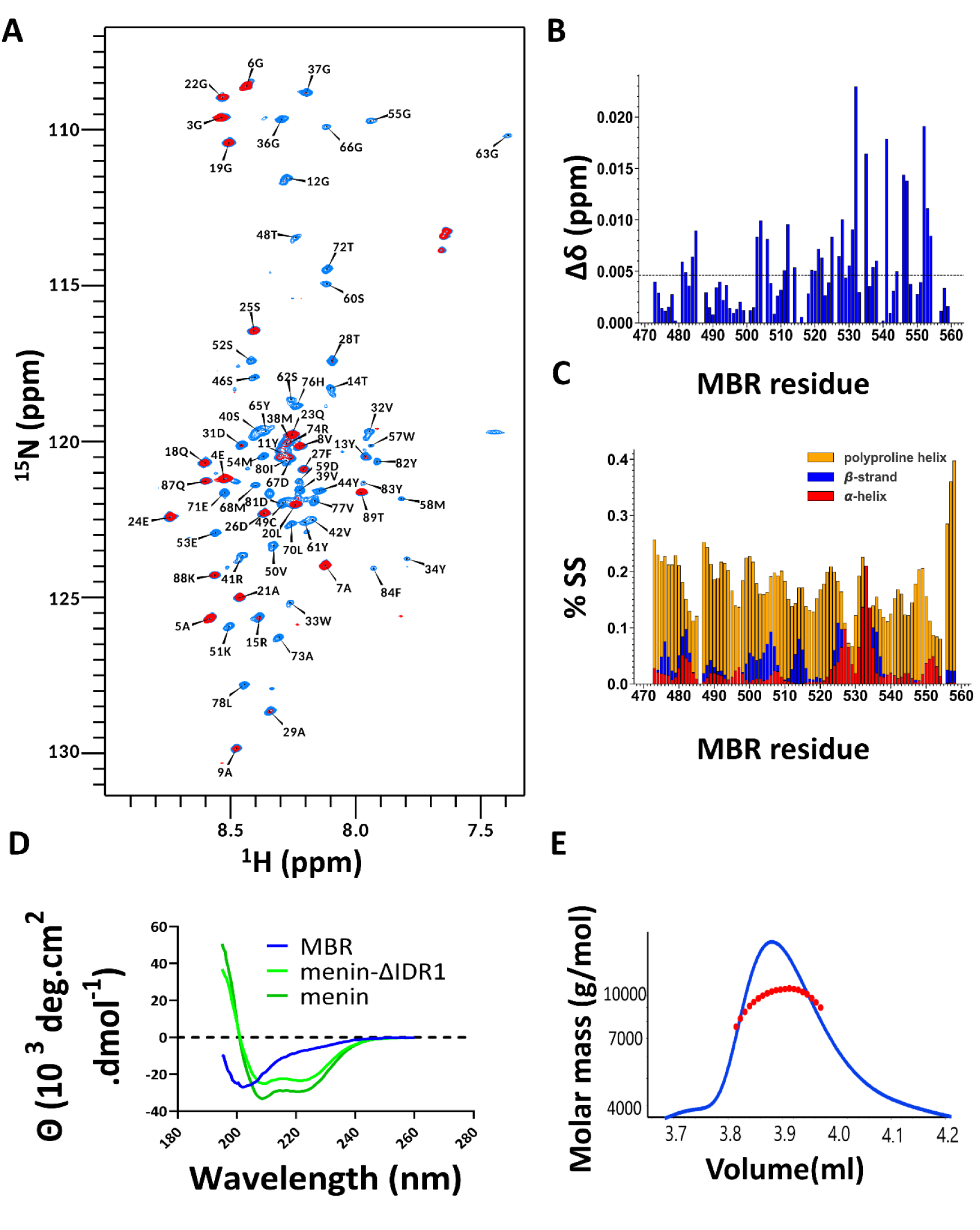
Menin interact with MBR. A) ^1^H ^15^N HSQC spectrum of MBR in the absence (blue) and presence (red) of menin B) Chemical shift perturbation (δΔ) between 100 μM MBR in the absence and presence of 130 μM menin. The dotted line shows the standard deviation in δΔ. C) Secondary-structure propensity within MBR as calculated from backbone chemical shift values using the δ2D server (https://www-cohsoftware.ch.cam.ac.uk/index.php/d2D): β-strand is shown in blue, PPII helix in yellow and ɑ-helix in red. D) CD spectra in the far-UV region 190-260 nm of MBR (blue), menin (green) and menin ΔIDR1 (green). Measurements were acquired at 25 °C with 2 μM menin/ menin ΔIDR1 and AR MBR at 7 μM concentration. E) SEC-MALS of MBR at 10 mg/ml in 50 mM Tris, 150 mM NaCl, 0.5 mM TCEP at pH 7.5.

We first assigned the backbone resonances of AR MBR as described in Materials and methods (Fig. 2A, Suppl. Table S1). To precisely delineate the menin binding site within AR MBR, next we have titrated AR MBR with the menin ΔIDR1, and determined characteristic chemical shift perturbations. In agreement with the above binding measurements with the two halves of AR MBR (MBR-A and MBR-B), there is a characteristic pattern of chemical shift changes within AR MBR (Fig. 2B, 1D, Suppl. Fig. S2B): larger chemical shift changes are evident in the C-terminal region (residues P534-F554), while smaller, still significant chemical shift perturbations occur within its N-terminus (Y481 and R485). This data confirm that MBR-B is the primary menin-binding region, whereas MBR-A contributes secondary, weaker binding (Fig. 1D,E, 2A, B).

Based on chemical shift information, we used the algorithm δ2D [27] to assess the secondary-structure preferences of AR MBR (Fig. 2C). These, in agreement with circular dichroism (CD) measurements (Fig. 2D), showed that the fragment consists of some transient structures, mainly of PPII helix, β-sheet, and a small percentage of α-helix (Fig. 2C). Deconvolution of the CD spectrum by BeStSel [28] showed a largely disordered structure, with 38% β-sheet, 56% coil and 5.8% α-helix content. This spectrum is in stark contrast with those of menin and menin ΔIDR1 (Fig. 2D), with dominant α-helix content (menin: 51% α-helix, 4% β-sheet, 45% coil, and menin ΔIDR1: 40% α-helix, 16% β-sheet, 44% coil). To underscore further the importance of menin-AR interaction, the stability of menin in thermal denaturation experiments was determined (menin and menin ΔIDR1 gave melting temperatures of 41.0 °C and 40.6 °C, respectively, cf. Fig. S2C). This is a low melting temperature for a human protein [29], thus the interaction and stabilization by binding partners like MLL and AR might be of prime importance for menin function. As menin tends to dimerize (see later) it is of interest to find out if AR MBR has any tendency to form dimers. To address this, SEC-MALS experiments were carried out. The apparent molar mass calculated was 9.5 kDa, which corresponds to the monomer of MBR (Fig. 2E). Cross-linking MBR and menin (Suppl. Fig. S3B, F,G) not only confirmed a direct interaction between the two proteins it already highlights that they might engage in the formation of higher-order complexes which may be relevant with their tendency to co-phase-separate.

### 3) Stoichiometry of menin-AR-MLL interactions

The IUPred score of menin shows that the C-terminus of menin is disordered (Fig. 3A). Menin appears as a monomer in the Protein data Bank (PDB 3u84) [10] and by AlphaFold [30] prediction (AF-A0A024R5D2) (Fig. 3B). Its domain organization suggests a large N-terminal domain, THUMB, and Palm (M1-V460), and a C-terminal region of two IDRs (R460-P519 and A586-L615, corresponding to a Finger domain). Different constructs of menin including menin ΔIDR1, menin ΔIDR1, IDR2 were used in the study (Fig. 3C). As our preliminary observations suggest that menin may form dimers, we thought to address its oligomeric state, as it may have far-reaching implications on the menin-AR-MLL interaction, suggesting a different regulatory model from the one described before [5]. We reasoned that if menin has different binding sites for MLL and AR, its potential dimeric nature will result in different competition or cross-linking scenarios if MLL and AR bind the same (overlapping) or different regions (Fig. 3D - F).

**Figure 3.**
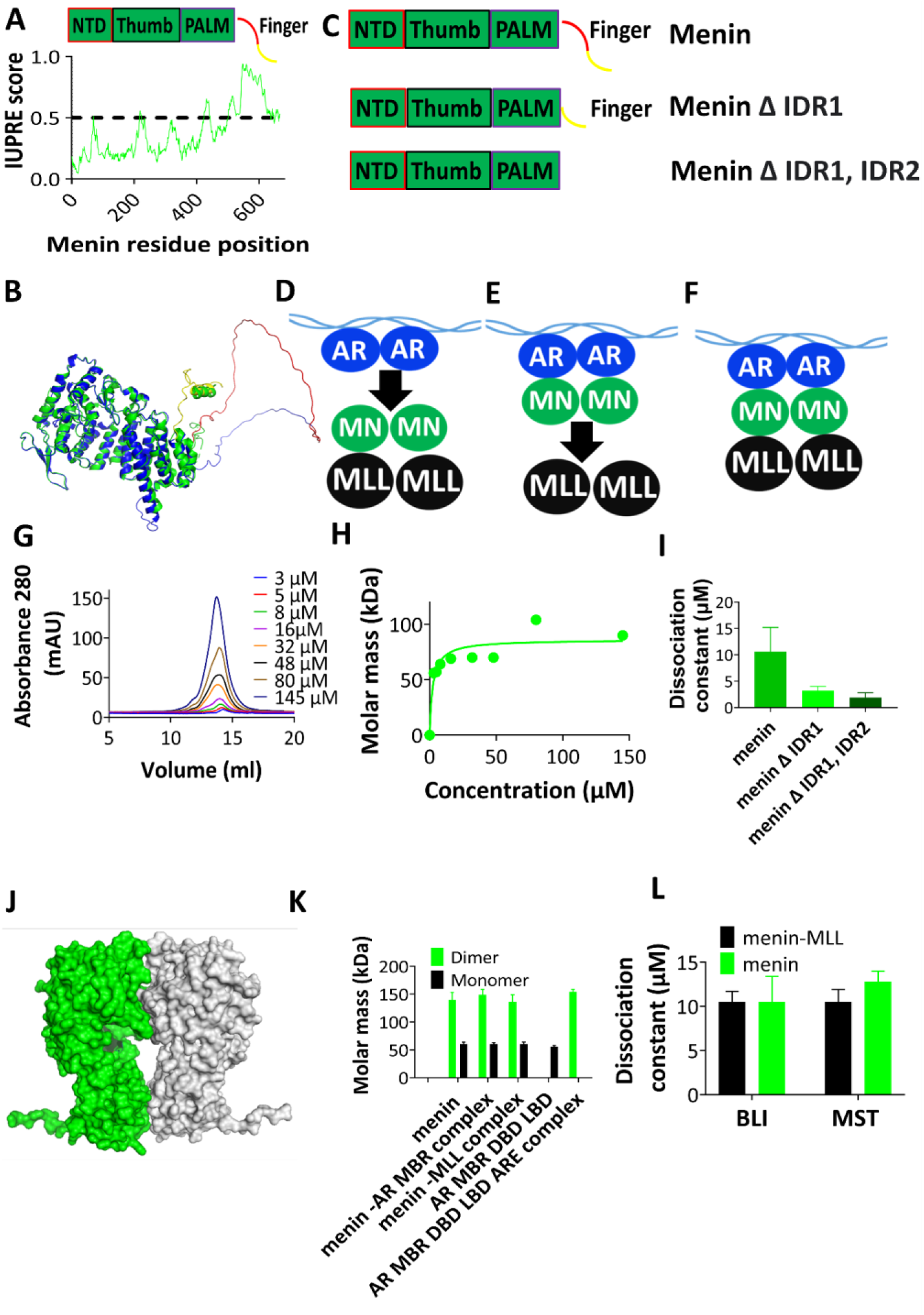
Possible stoichiometry and organization of the menin-AR-MLL complex. A) The organization of four regions of menin, the N-terminal ordered domain (NTD), Thumb, and PALM, and the intrinsically disordered C-terminal Finger domain.IUPred prediction for disorder propensity of menin. B) Crystal structure of menin (PDB 3u84, green), superimposed on the structure predicted by AlphaFold (AF-A0A024R5D2, blue). Two menin deletion constructs have been used in this study, the menin ΔIDR1 has region R460-P519 deleted, whereas the menin ΔIDR1, IDR2. has both R460-P519 and Q586-L615 deleted. C) Representation of different constructs of menin used, with two deletion constructs including menin ΔIDR1 and menin ΔIDR1, IDR2. D,E,F) Possible models of AR-menin-MLL interaction, assuming overlapping MLL- and AR-binding sites on menin, enabling alternative menin-MLL and AR-menin complexes (D, E) or different interaction sites enabling the formation of a ternary complex between dimeric (ARE-bound) AR, dimeric menin and two MLL subunits (E). G) Gel filtration of menin at different concentrations ranging from 3 µM to 145 µM (using an S200 10/300 column). H) From the elution volumes of gel-filtration (G), the apparent molar mass of menin is calculated by calibrating the column with a BioRad calibration standard (cf. Suppl. Fig. S3A). I) Dissociation constant, Kd, of menin dimerization, in the case of different constructs (as defined on panel C). J) Structure of dimeric menin (monomer, PDB 3u85) predicted by PDBePISA [31] K) Dynamic light scattering (DLS) of menin, to determine its molar mass in the presence and absence of MLL and AR as well as molar mass of MBR-DBD-LBD and MBR-DBD-LBD complex with ARE. L) Dissociation constant (Kd) of the interaction of menin, and the menin-MLL complex, with AR

First, we used gel filtration (size-exclusion chromatography) at different menin concentrations (3 to 145 μM) to determine its apparent Mw (Fig. 3G). We found that the apparent Mw shifts from 56 kDa (at 3 μM) to about 104 kDa (at above 48 μM, cf. Fig. 3H). Considering that menin is a 61-kDa protein, this shift in its observed Mw suggests its transition from a monomer to a dimer by increasing concentrations.To further confirm the dimeric nature of menin, we have also carried out cross-linking experiments with glutaraldehyde, which showed the formation of dimer and higher-order oligomeric products (Suppl. Fig. S3B).

To further confirm the dimeric nature of menin, SEC-MALS was used for menin, menin ΔIDR1, and menin ΔIDR1, Δ IDR2. The apparent Mw calculated for the menin ΔIDR1 at 60 μM is 86 kDa (Fig. 2H), for menin at 60 μM and 350 μM is 103 kDa and 164.3 kDa, respectively (Suppl. Fig. 4A, B, C). In case of menin ΔIDR1, IDR2, its apparent Mw at 30 μM is 65.4 kDa, at 130 μM it is 87.1 kDa, whereas at 390 μM it is 102.0 kDa. These results confirm that menin undergoes concentration-dependent dimerization.

We have also performed high-performance liquid chromatography - small angle X-ray scattering (HPLC-SAXS) experiment. The change in the radius of gyration (Rg) for each frame (Suppl. Fig. S4D) suggests a molar mass of 117.7 kDa, corresponding to a dimer.

As the above SEC-MALS experiments suggested that the disordered C-terminal domain of menin is involved in dimerization, we wanted to further address this question by measuring the affinity of monomer-monomer interaction by MST. We observed Kd = 10.4 ± 4 μM for menin, Kd = 1.9 ± 0.9 μM for menin ΔIDR1, and Kd = 3.2 ±0.8 μM for menin ΔIDR1, IDR2(cf. Suppl. Fig. S3 C, D,E), which suggests that the dimeric interface is on the N-terminus of the protein and the C-terminal IDRs affects menin dimerization. To enable a low-resolution structural model of the system, we wanted to know the relative orientation of the subunits in the dimer, i.e., where the dimeric interface is located within menin, especially if the disordered C-terminal IDR is involved. Based on this information (Fig. 3I, cf. Suppl. Fig. S3 C,D,E), we suggest a model for the structure of the menin dimer, as predicted by the PDBePISA algorithm [31] based on the monomeric PDB structure (PDB: 3u85, cf. Fig. 3J). Another PDBePISA algorithm, alpha fold was also used for prediction of dimeric interface (Suppl. Fig. S E-G). In all these models we majorly have one region in the N terminus of menin which is important for maintaining the dimer interface.

To confirm the image provided by Rg, we also asked the radius of hydration (Rh) by dynamic light scattering (DLS). The Rh of monomeric menin is 3.3 nm with a corresponding molar mass of 60 kDa, whereas for dimeric menin it is 5 nm, with a corresponding molar mass of 125 kDa (Fig. 3K). Next we wanted to know whether ARE has any influence on AR MBR DBD LBD dimerization, to know that first we measured the mass of AR MBR DBD LBD without ARE and it was 56 kDa and with ARE it was 154 kDa. This means that ARE triggers the formation of AR dimer. As AR DBD forms a dimer upon binding to ARE [19], and there might be a functional interaction between AR and menin, the dimeric nature of menin may have a basic influence on AR DNA binding (cf. Fig. 1C-E). Thus, we next asked if MLL and AR, two binding partners, have any influence on the oligomeric state of menin. Upon adding five times the concentration of AR MBR and MLL to menin, no significant difference in its hydrodynamic radius and molar mass was observed, i.e., the binding of AR MBR or MLL does not affect the oligomerization state of menin, but the complex of AR MBR-DBD-LBD with ARE does trigger the dimerisation of menin (Fig. 3K, 1D).

A further critical question with regards to the regulatory circuit made up of these proteins is whether AR and MLL have different or the same (overlapping) binding sites on menin, as these may enable only alternative binding of AR and MLL to menin (Fig. 3D, E), or the simultaneous binding of both proteins, enabling the assembly of a ternary AR-menin-MLL complex (as depicted on Fig. 2F). To distinguish between these possible modes of organization, we used MST and BLI experiments. We found that menin interacts with AR MBR with a Kd = 10.5 ± 0.9 μM (BLI), and 9.8 ± 2.0 μM (MST), whereas the menin-MLL complex interacts with MBR with a Kd of 10.5 ± 0.9 μM and 10.8 ± 0.64 μM, respectively (Fig. 3L). As the interaction of the menin and menin-MLL complex with AR are the same to within experimental error, we may conclude that AR and MLL apparently have different binding sites on menin, i.e., an AR-menin-MLL ternary complex as depicted on Fig. 3F can form.

### 4) Menin-binding motif of MLL interacts with RNA to drive phase separation

It has been shown that menin interacts with the N-terminal region of MLL1 and MLL2. MLL1/2 is a scaffold of a large complex composed of more than 30 units e.g ASH2L, PbBP5, WDR5 and these components are important for Histone methyltransferase activity [32]. In this work we study MLL1 (and refer to it as MLL). As menin interacts with the N terminus of MLL, it is present in complexes assembled by wild-type MLL and its fusion proteins [4]. The interaction region of menin and MLL has been narrowed down to the region 1 M-40 L, which has two menin binding motifs (MBMs), underscored by X-ray crystallography, NMR and other biophysical techniques [9,10]: the two MBMs are referred to as MBM1 and MBM2 [10]. The domain architecture of MLL (Fig. 4A) shows excessive structural disorder throughout the entire sequence. As MBM1 and MBM2 are both localized within the first 40 residues of MLL, we generated four constructs to study the consequences of the interaction they drive (Fig. 4B): MLL_Pep1 contains MBM1 and MBM2, MLL_Pep2 contains MBM1, MLL_Pep3 is an MLL_Pep2 mutant in which the two terminal arginines are mutated to glycines, while MLL_Pep4 contains an oligo-glycine stretch present in MLL [10].

**Figure 4.**
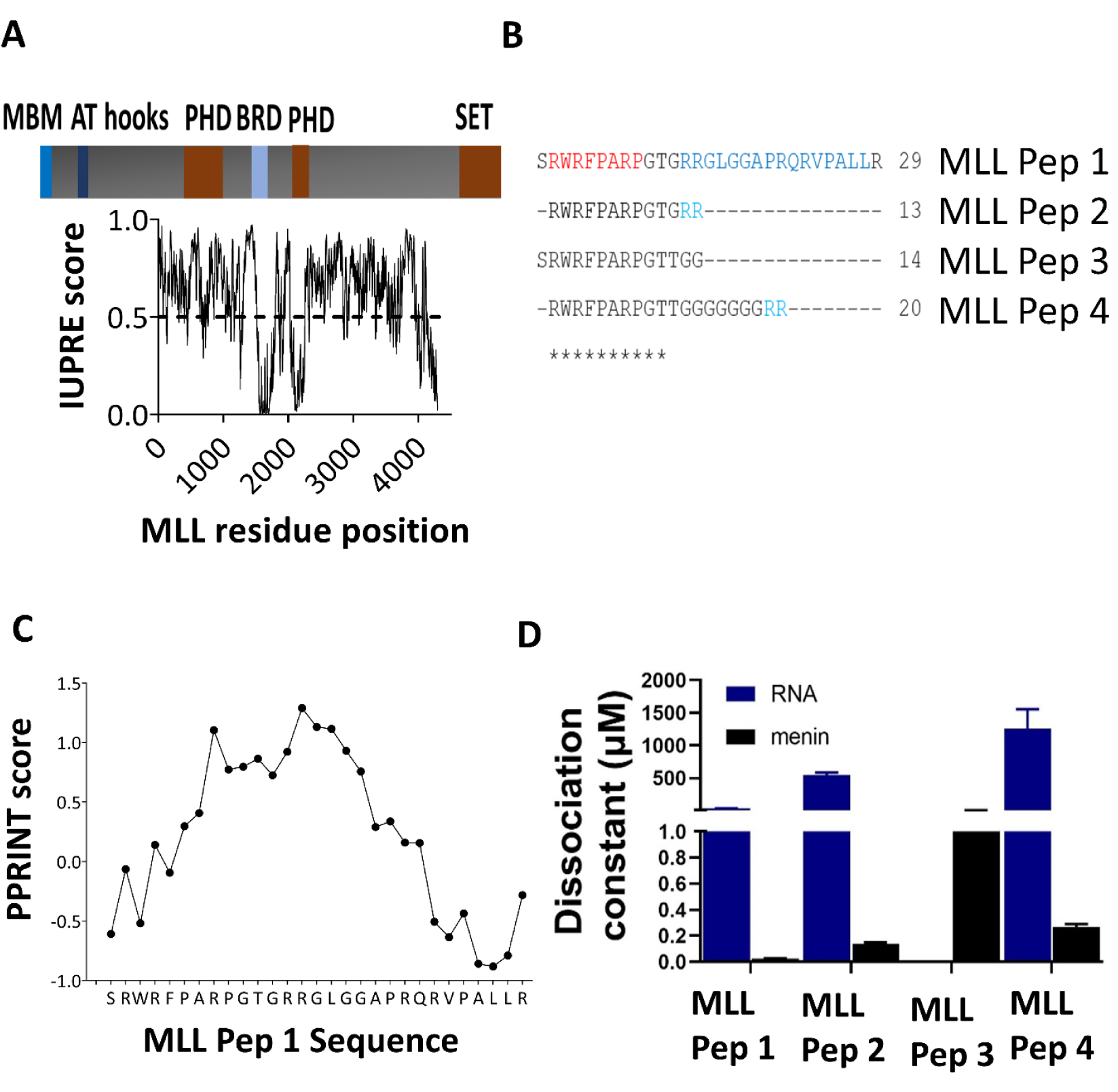
MLL interacts with RNA. A) MLL is a protein of 3969 amino acids, with a menin-binding motif (MBM) located at the N terminus, followed by an AT-hook, PHD, BRD (Bromo-), and SET domains in a largely disordered protein, as demonstrated by its IUPred score. B) MLL Peptides Pep 1 through Pep 4, synthesized for menin- and RNA-binding studies, encompassing the two menin-binding motifs MBM 1 (red) and MBM 2 (blue) C) PPRINT score predicts RNA-binding propensity within MBM [33]. D) Interaction of menin and RNA (poly U) with different MLL peptides (defined in Panel B).

By applying the PPRINT predictor [33], we could ascertain that MBM1 may also carry an RNA-binding motif (Fig. 4C), which has not been experimentally proven, although it could basically regulate the menin-MLL interaction. First we used MST to demonstrate MLL-RNA interaction and we found that MLL_Pep1 interacts with RNA (PolyU) with a Kd = 31.18 ±1.6 μM, MLL_Pep2 with Kd = 548 ± 36 μM, whereas MLL_Pep3 did not bind and MLL_Pep4 binds very weak, with a Kd = 1254 ± 301 μM only. From these differences, we can conclude that the two arginines within Peptide 1 (R24, R25 highlighted in blue on Fig. 3B) are important for interaction with RNA. We also assessed the interaction of MLL Peptides with menin, which gave Kds = 24.7± 3 nM (MLL_Pep1), 140 ± 11 nM (MLL_Pep2), 31.3 ± 0.5 µM (MLL_Pep3), and 268.7 ± 23 nM (MLL_Pep4). From these values, we conclude that the MBM region in MLL interacts with both menin and RNA, and in both interactions both MBM1 and MBM2 are important, with the R13R14 motif playing a key role (Fig. 4D, Suppl. Fig. S5 A-G).

We next observed that the interaction of this RNA-binding region with RNA drives liquid-liquid phase separation (LLPS) of MLL, as first shown by increasing turbidity at OD 600 for MLL Pep 1 in the presence of Poly U (Fig. 5A), confirmed by microscopic images of droplets formed (Fig. 5B). Fluorescence recovery after photobleaching (FRAP) showed that the droplets are very dynamic, recovering with a half-life of about 10s (Fig. 5C). A similar trend was observed for MLL Pep 2 (Fig. 5D, E, F). The importance of RNA binding for LLPS is shown by the fact that MLL_Pep3, which lacks the two arginine that mediate the interaction with RNA, undergoes no LLPS (Fig. 5G, H). MLL_Pep4 formed condensates with similar dynamics (Fig. 5I-K).

**Figure 5.**
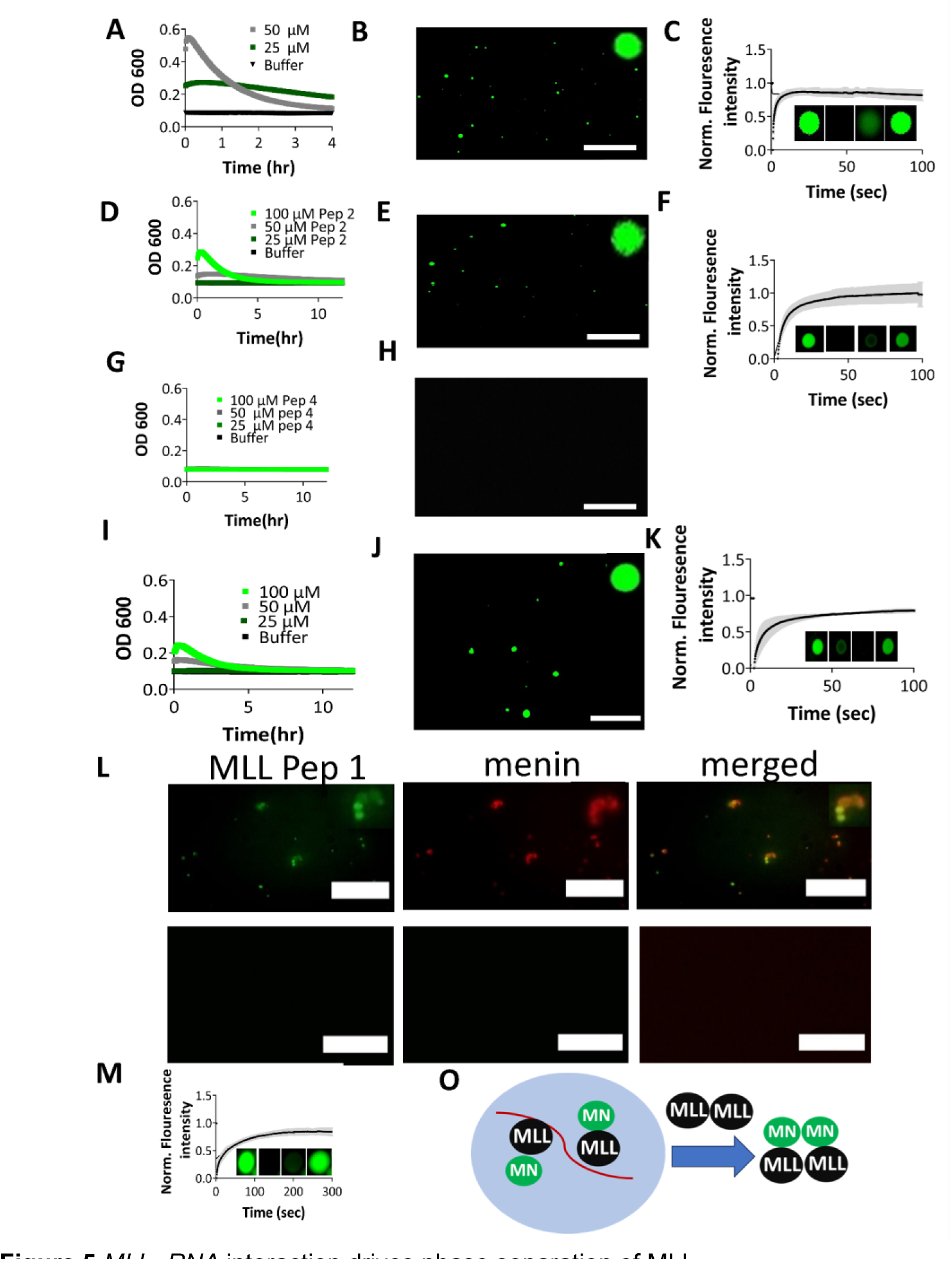
*MLL -RNA* interaction drives phase separation of MLL. (A-K) LLPS of peptides (at 25 μM) with 0.5 μg/μl RNA (polyU) approached by OD 600, microscopy, and FRAP for MLL Pep 1 (A, B, C), Peptide 2 (D, E, F), Peptide 3 (G, H), and Peptide 4 (I, J, K) L) Microscopic images of MLL droplets in the presence of low (10 µM) and high (250 µM) concentration of menin M) FRAP of MLL_Pep1 in the presence of menin at 10μM. O) model for regulation of MLL phase separation by menin. Scale bars in all microscopic images are 20 μm.

As RNA- and menin-binding regions of MLL overlap, we next asked if menin influences the condensation driven by MLL-RNA interaction. We found that MLL-RNA droplets still form in the presence of menin, and menin gets recruited into MLL-RNA condensates, with a tendency to localize to droplets, promoting the clustering of droplets (Fig. 5L, Suppl. Fig. S5 H). Further, menin makes the droplets less dynamic, as their complete FRAP recovery requires about 300s, in comparison to 10s in the absence of menin (Fig. 5M vs. Fig. 5C). Although the menin-MLL interaction is, in principle, much stronger than that of RNA-MLL, this result suggests that menin does not compete with the MLL-RNA interaction, rather its “coats” the droplets, making them more isolated form solution, thus hindering diffusion in and out of droplets. Invasion of droplets does occur, though, at higher menin concentrations, menin has an inhibitory effect on MLL-RNA LLPS (Fig. 5L, M,O).

Overall, these results show that the N terminus of MLL can interact with RNA, promoting its LLPS, leading to the formation of condensates. Menin is important for regulating these condensates, and either solidifies condensates or completely inhibits phase separation, in a concentration-dependent manner (Fig. 5L, M, O). Given the well-defined region within MLL that mediates these interactions, these findings may also point toward potential specific targeting of the interaction of MLL with RNA and/or menin in developing cancer drugs [7].

### 5) Menin is also recruited into condensates formed by androgen receptor

AR is known to form condensates promoted by RNA [12,13]. In our previous work, we showed that the DNA-binding domain (DBD) of AR (cf. Fig. 1C) drives AR phase separation [12]. Here, to explore details of these effects further around the DBD, we generated different constructs of AR (Fig. 1). As DBD and LBD are connected by a short disordered “hinge”, and a special feature of AR-v7 is that it carries here an alternative sequence, “alt hinge”. We generated a range of constructs: removed LBD, varied the hinge region, and also included or deleted the disordered menin-binding region (MBR) N-terminal to DBD (Fig. 4A). We then first addressed the RNA-binding of these various constructs with the ARE DNA, resulting in Kd = 0.2 ± 0.01 μM for AR DBD-hinge, 0.71 ± 0.05 μM for AR DBD-alt hinge, and 1500 ± 20 μM for MBR-DBD. Apparently, regions flanking DBD allosterically regulate the interaction of DBD with ARE, the hinge region positively, whereas MBR negatively regulates the interaction (Suppl. Fig. S6 A). Another important question is whether AR is present as a monomer or dimer in solution, as it is related to the question of the mode of AR-menin-MLL assembly (cf. Fig. 3D-F, K). To answer this question, we applied SEC-MALS, to measure the apparent Mw of DBD (11.6 kDa at 1 mg/ml, and 12 kDa at 5 mg/ml), which suggests it is a monomer (its absolute Mw is 9.8 kDa). For the AR DBD-hinge construct, its molar mass obtained (13.2 kDa at 1 mg/ml and 14.0 kDa at 5 mg/ml, close to its absolute Mw, 15.6 kDa) also suggests a predominantly monomeric state in solution (Fig. 4C, D). As the hinge region exerts very strong positive allostery on DNA binding, it could be due to promoting DBD dimerization, as DBD of AR is a dimer when it binds ARE(cf. Fig. 3K) [19].

Next, we explored the phase-separation behavior of these proteins, and asked whether the binding affinity of constructs with RNA has any effect on their phase-separation tendency. First Dynamic light scattering (DLS) was applied to DBD-alt and MBR-DBD-alt constructs and Rh for DBD-alt plateau at 1000 nm and for MBR-DBD-alt at 1500 nm, presence of MBR increased the size of droplets to 500 nm in presence of RNA (Suppl. Fig. S6D).

Next a very important question is that phase separation behavior of AR vs AR-v7 Because of the presence of the ligand-binding domain (LBD) in full-length (FL) AR, we added dihydrotestosterone (DHT) to bring the protein to its active conformation, characterized by an intramolecular interaction between its N- and C-termini [34], when we observed the formation of condensates upon the addition of RNA, not seen previously *in vitro*. These condensates are dynamic, as they recover by FRAP to 90 % in 100s (Fig. 6A). Next, we tested the behavior of AR-v7 in the presence of RNA. AR-v7 forms small clusters, which are very different from that of AR in shape and size as well as recovery, characterized by a faster rate (10s vs. 40s) of recovery in FRAP (Fig. 6B). These results point to the basic influence of LBD on phase separation of AR and may suggest the opportunity of specific targeting of clusters of AR-v7 as compared to FL AR (Fig. 6A, B).

**Figure 6.**
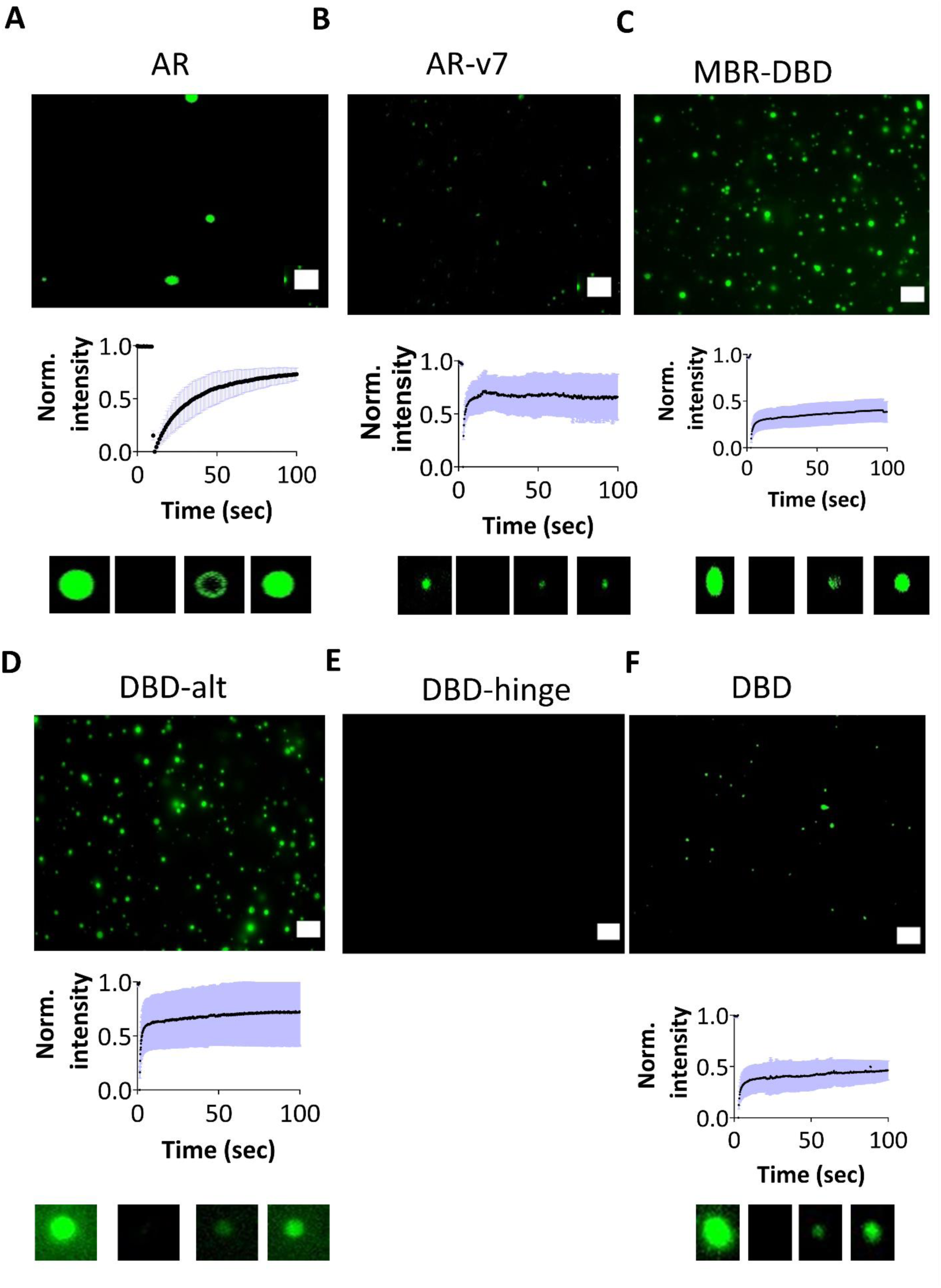
AR phase separate differently than pathological variant, AR-v7. A) Full-length (FL) AR undergoes condensation in presence of DHT and 5 mM ATP, B) AR-v7 clusters. Condensates of C) MBR-DBD, D) DBD-alt hinge, E) DBD-hinge, F) DBD at 50 μM. All microscopic images were acquired at 0.25 μg/μl polyU; FRAP shows recovery of their fluorescence.

By analyzing the influence of various domains flanking the DBD on LLPS, we made the interesting observations that their effect is the opposite to that made on DNA binding: while MBR-DBD, DBD-alt hinge, and DBD itself form condensates, DBD-hinge does not (Fig. 6 B,C,D,E,F). DBD without any flanking region forms small clusters, because of its weaker interaction with RNA [12]. It is very interesting that DBD-hinge does not form condenses, suggesting a basic regulatory role of the hinge on DNA/RNA binding and phase separation of the protein (Fig. 6E). The FRAP recovery shows that different regions of AR are dynamic in nature and intensity is recovered to more than 50% after photobleaching (Fig. 6). Whereas these results were obtained with an RNA mimic (polyU), it was also shown that total RNA extracted from LNCAP cell lines triggered phase separation and affected the phase separation behavior of AR in a similar manner (Fig. 7A).

**Figure 7.**
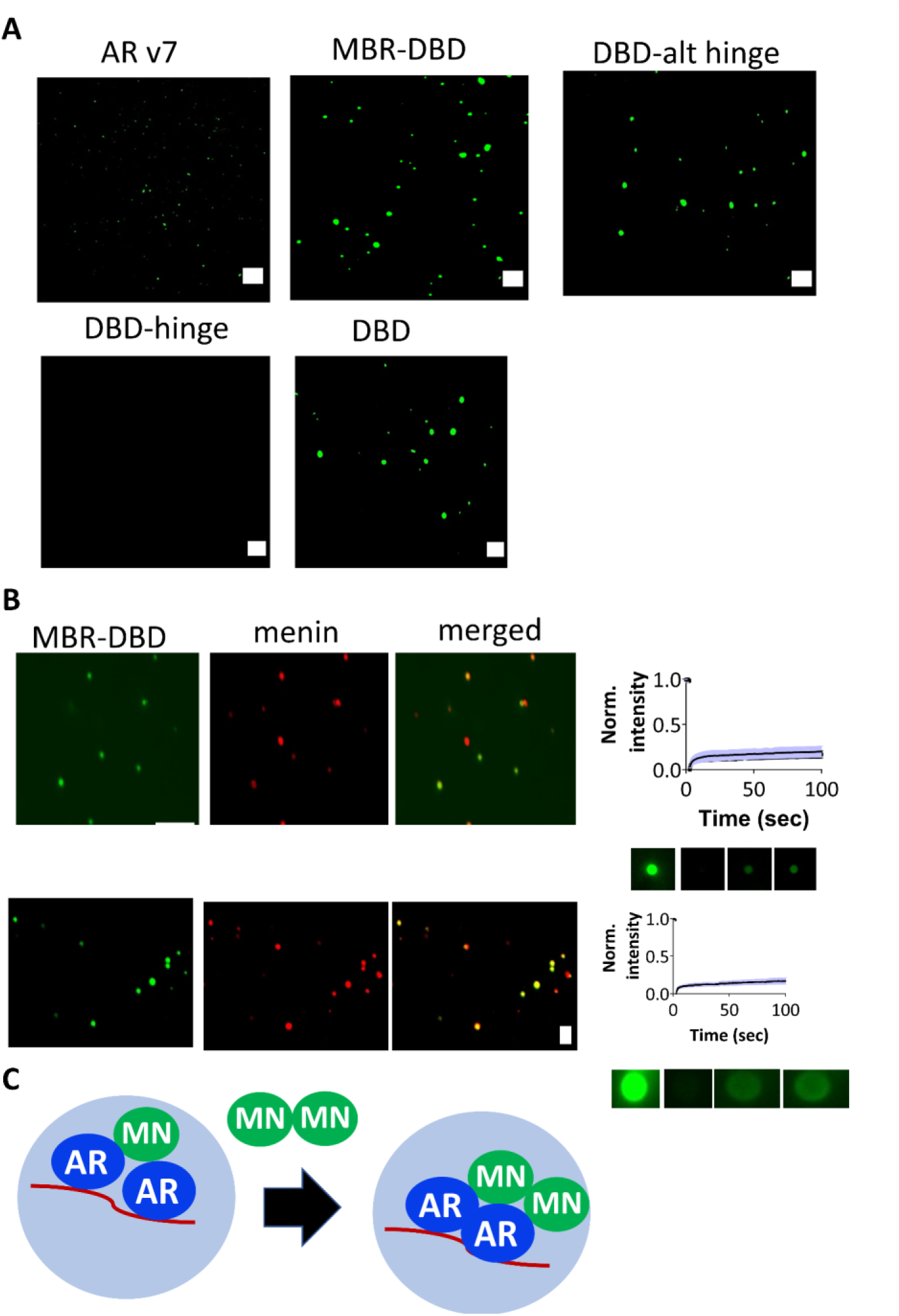
Menin is recruited into AR condensates. A) microscopy of droplets of different AR constructs, with total RNA (0.25 mg/ml) extracts from LNCAP cells. B) microscopy of droplets of MBR-DBD (25 μM) at two different menin concentrations, 10 μM (upper panels) and 250 μM (lower panels), along with their FRAP curves. C) A model for phase separation of AR and recruitment of menin, suggesting that menin at high, but not at low, concentration, menin is exclusively localized into AR droplets. Scale bars on microscopic images, 10 μm. The buffer used in the experiments is 50 mM phosphate, 75 mM NaCl, pH 7.0, TCEP 0.5 mM TCEP.

Next, we asked the effect of menin on the phase separation of AR, with the MBR-DBD construct that carries the menin-binding region of AR, and promotes the LLPS of DBD. We found that at a low (10 µM) concentration of menin, some, but not all, AR droplets became enriched with menin, whereas at a high (250 µM) concentration, menin co-localizes with AR while the droplets turn to more gel-like, as shown by their FRAP profile (Fig. 7B).

Overall, these experiments show that FL AR forms condensates, whereas AR-v7 tends to form smaller clusters. The flanking regions of DBD affect phase separation mediated by DBD, whereas AR droplets recruit menin and change their dynamics, leading to the formation of gel-like states.

### 6) The menin-AR-MLL complex forms liquid condensates in cells

Next, interactions and phase separation events observed *in vitro* also express themselves inside living cells. For this purpose, we used LNCAP, HEK, and U20S cell lines, into which GFP-fused AR (GFP-AR) was transiently transfected and stimulated by DHT. We found that upon ligand stimulation, AR translocate to the nucleus and forms foci, as already reported [13,35]. The AR foci are present in the whole nucleus except the nucleolus, as seen on fluorescence intensity plotted across the nucleus of all three cell types (Fig. 8 A, B, C). To probe the liquid nature of these condensates, we applied 1,6 hexanediol (2.5%), a compound known to inhibit weak hydrophobic interactions and interfere with LLPS, both *in vitro* and in cells [36,37]. 1,6 hexanediol results in inhibition of the formation of AR foci in all three types of cells (Fig. 8 A-C).

**Figure 8.**
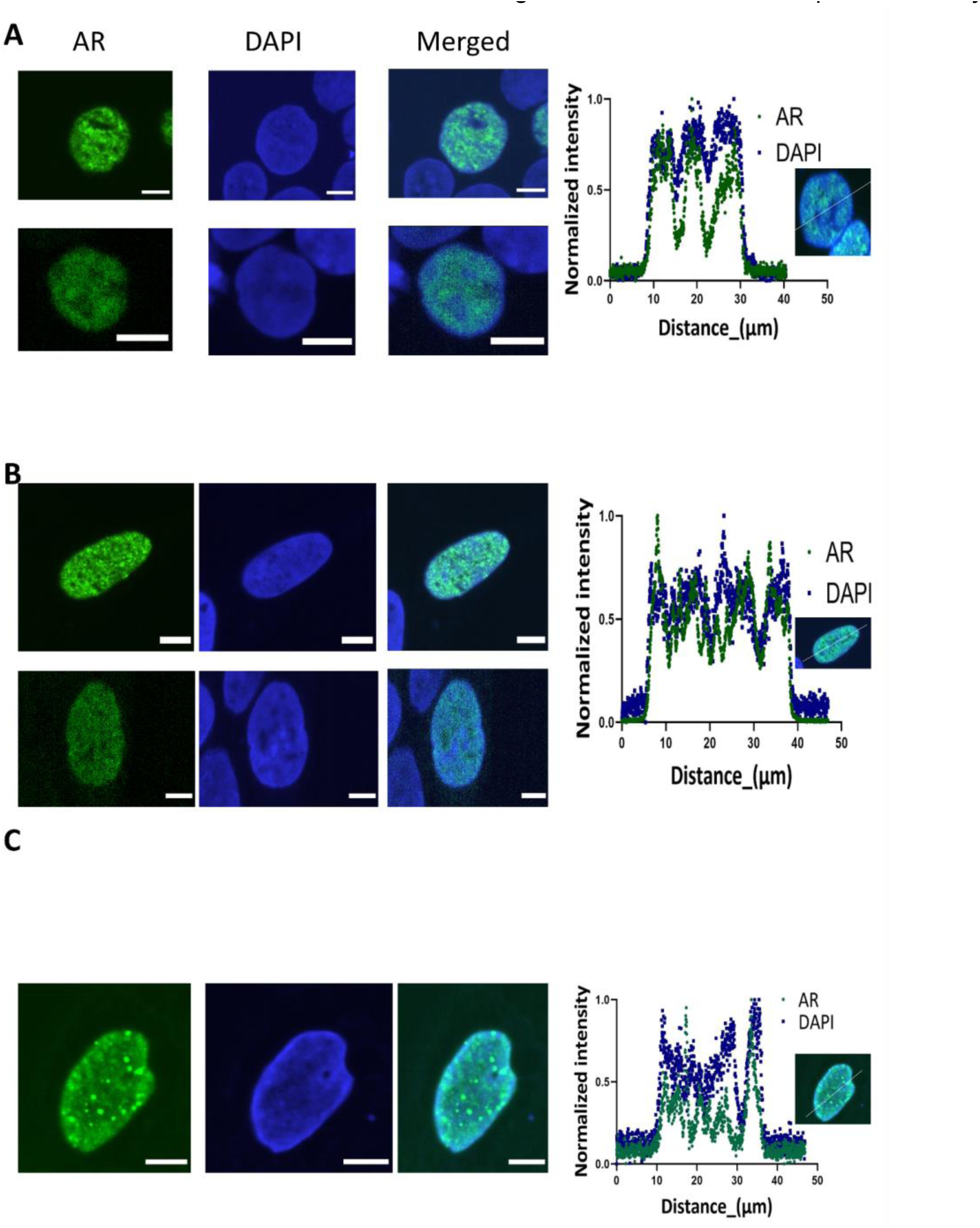
AR condensates in cell. (A, B, C) HEK, LNCAP and U20S cells, respectively, expressing GFP-AR in the cell nucleus. Cells were starved with 10% charcoal-stripped FBS (CS-FBS) media for two days to cause hormone deprivation and stimulation was performed by 10 nM DHT with and without 2.5 % hexanediol (Hexa) for 15 minutes. On the right panels, fluorescence intensity scans across the whole cell are shown, underscoring the formation of positive foci in the nucleus, but not in the nucleolus.

The next question we asked is if menin and MLL are recruited into AR condensates as suggested by our prior *in vitro* results. In HEK and LNCAP cells expressing GFP-AR, the recruitment of menin and MLL was observed by immunostaining with respective primary antibodies and fluorescently conjugated secondary antibodies (Fig. 9A-C). In HEK cells, AR expression, and stimulation by DHT resulted in the simultaneous recruitment of menin and MLL, to form the AR-menin-MLL complex. When AR is not expressed, menin and MLL co-localize in the nucleus, MLL forming small clusters and menin forming bigger foci. In the presence of AR, the intensity measured across cells shows the colocalization of three binding partners: whereas menin and MLL are expressed in both the nucleus and cytoplasm (Fig. 9A, B, C), there is a significant difference in the recruitment of MLL in the presence and absence of AR (Fig. 9D) while there is no difference in the recruitment of menin.

**Figure 9.**
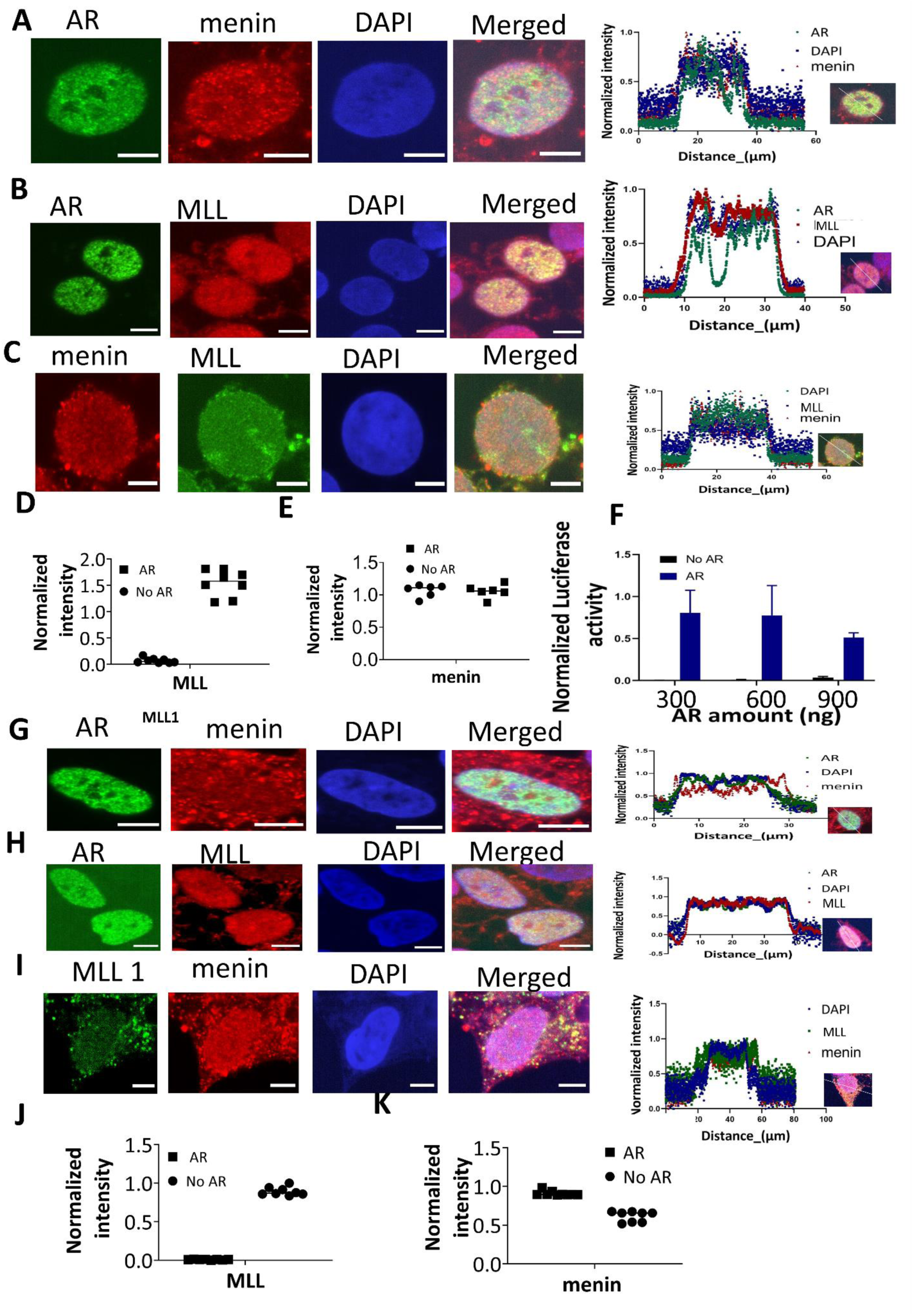
Complex menin-AR-MLL condensates in cell. Confocal images of cells starved with 10 % charcoal-stripped FBS (CS-FBS) media for two days to cause hormone deprivation, enabling an effective stimulation by 10 nM DHT, of: (A) HEK cells expressing GFP-AR in the cell nucleus, menin is stained with primary anti-menin (1:1000 dilution) and secondary cy5 conjugated anti-mouse (1:1000) antibody, (B) primary anti-MLL (1:500) and secondary cy5-conjugated anti-rabbit (1:1000) antibody, (C) anti-menin (1:1000 dilution) and anti-MLL (1:500) antibody, followed by secondary anti-rabbit (1:500) and anti-mouse (1:1000 dilution) antibodies. Intensity profiles for AR, menin, MLL, and DAPI along a white straight line crossing the nucleus of representative cells were obtained by the image j software. Normalized intensity of MLL (D), and menin (E) with and without AR expression, is shown. F) AR activation in HEK 293T cells transfected with FL AR and pARE-E1b-Luc. GFP-tagged AR expressed in LNCAP cells and immunostained for: menin (G, red), and MLL (H, red). I) MLL (green) and menin (red) without expressing FL AR. Intensity profiles of AR, menin, MLL, and DAPI for LNCAP were obtained along a white straight line crossing the nucleus of representative cells by the image j software. J) Intensity profile of MLL, and menin (K) compared with and without AR expression

Next, we addressed the effect of AR expression and recruitment on the menin-MLL interaction, and whether this recruitment of MLL influences transcription and translation. For this purpose, a luciferase assay using two plasmids was adapted: one plasmid expressing AR and the second expressing luciferase reporter, downstream of ARE. Co-transfection and stimulation by DHT result in an increase in cellular luciferase level, compared to control (Fig. 9F). Next, the same set of experiments was performed in LNCAP cells. Upon AR transfection and stimulation by DHT, the recruitment of menin and MLL changed significantly, showing colocalization of three components (Fig. 9G-J) in the foci formed, suggesting the underlying menin-AR-MLL interaction, a behavior very different from recruitment in HEK cells.

In both cell lines, menin and MLL, unlike AR, are also expressed in the nucleolus. Previously, not only MLL but its fusion proteins have also been reported to be enriched in the nucleolus [38] [39], where it colocalizes with menin. Because in its fusion proteins the N-terminus of MLL is retained, menin can bind both intact MLL and its fusion products (Fig. 9), driving the expression of different genes and recruitment of different partners resulting in prostate cancer. This data confirm that menin-AR-MLL form condensates in the cell lines studied, and expression of AR results in the recruitment of both menin and MLL, resulting in an increased transcriptional activity.

## Discussion

AR interacts with more than 200 binding partners [40], which trigger, repress or modulate its transcriptional activity. Some of these binding partners, such as Mediator 1 (MED1) [13,40,41], have been characterized for their recruitment to AR condensates and/or regulation of AR function, but the recruitment of most other binding partners to AR condensates, and their effect on the transcription activity of AR is unexplored [13,42]. Recently, it has been suggested that AR, menin and MLL form a transcription module that regulates the transcription activity of AR [5]. In order to characterize the readout of the underlying interactions, we have analyzed AR-menin and menin-MLL interactions in detail, as this module appears to be misregulated in cancer [5,43]. Our basic observation relevant with the topology of organization is that menin forms a dimer in a concentration-dependent manner, which gives rise to a variation of potential assembly of ternary complexes and its regulation by LLPS at two scenarios, AR overexpression and MLL overexpression (Fig. 10), which are realized in different cancer types.

**Figure 10.**
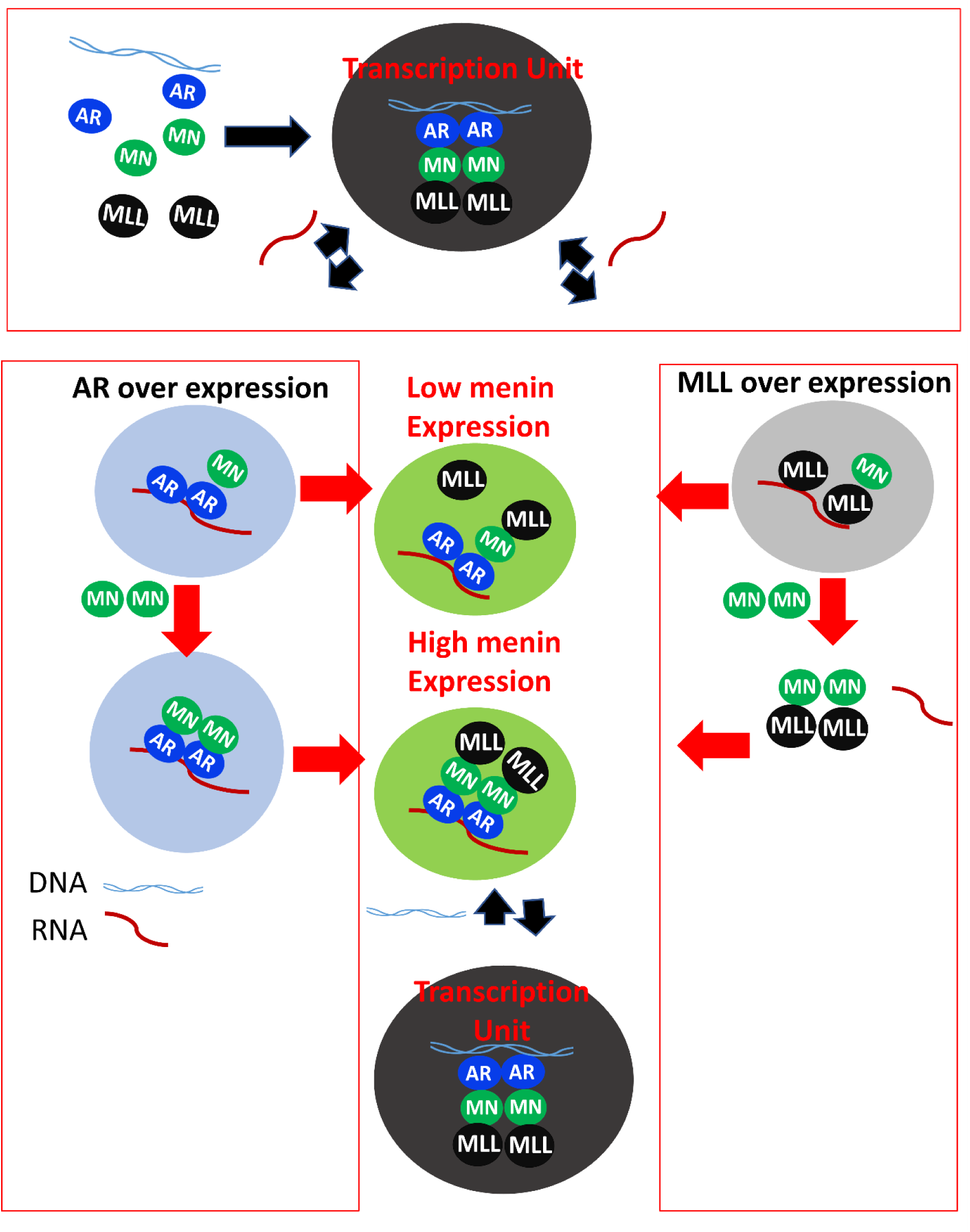
Suggested model of the molecular regulation of menin-AR-MLL complex. Menin is a dimer that has separate binding sites for AR and MLL, thus it can simultaneously bind to AR and MLL, resulting in a ternary complex of dimeric (DNA-bound) AR, dimeric menin and two subunits of MLL. This assembly is regulated by different factors, including DNA, RNA, and the actual concentration of menin. At low concentrations, menin is recruited into condensates of AR and MLL, whereas at high concentrations of menin, RNA is completed off from MLL and results in the inhibition of condensates. Condensates of AR recruit menin at a high concentration of menin and it turns more gel-like.

Within the menin-MLL-AR system, menin has specific binding regions on both AR (MBR, within its disordered NTD) and MLL (MBM, within its disordered N-terminal RNA-binding region). Binding to AR is regulated by a cooperative interplay between AR domains and regions, as AR DBD inhibits menin binding, whereas the activation function 1 (AF1) within its NTD, and the hinge specific to AR not its cancer-associated isoform AR-v7, counteract this inhibition. Interestingly, although a large fraction of AR is intrinsically disordered, allosteric interplay between remote regions of IDPs and IDRs is enabled by rearrangements in the structural ensemble, as captured by the ensemble view of allostery [43–45] that could results in very complex signaling switches regulated by multistery [45,46]. These mechanisms may result in both agonistic and antagonistic interplay between different regions, and such long-range communication within disordered regions of AR, and other nuclear hormone receptors, has already been observed [47].

Within MLL, menin binds via a binding motif within its RNA-binding region. Binding of at low menin to MLL does compete with that of RNA, enabling the buildup of larger ternary assemblies, as apparent from the recruitment of menin to MLL-RNA condensates but at high concentration menin does compete off RNA to inhibit LLPS. A similar logic applies to menin interacting with AR, as MBR and DBD are distinct regions of AR, enabling simultaneous AR-menin and RNA-AR interactions, which may be important in the formation of complex condensates. It is of note that the AR- and MLL-binding regions on menin also do not overlap, enabling the simultaneous binding of MLL and AR (and potentially, RNA) to menin [48] (Fig. 10), which may be important for co-phase separation of the entire quaternary system, which has the capacity to seed AR superenhancers [13,49]. We have dissected these relations by studying the LLPS of AR-RNA and MLL-RNA separately.

We found that all domain combinations of AR studied form condensates with RNA, but AR-v7 promotes different types of droplets than FL AR, as also implicated in the literature [35]. In addition, the native DBD-hinge has much less LLPS propensity than pathological DBD-alt hinge. These results confirm that AR-v7 has an aggravated tendency to phase-separate and be more dynamic. Interestingly, menin at both low and high concentrations (i.e., both in its monomeric and dimeric state) is recruited into these condensates, significantly reducing their dynamics, i.e., making them gel-like at high concentrations. MLL-RNA condensates also recruit menin, making these less dynamic at low concentrations, and inhibiting their formation all together at high concentrations.

These results of MLL-RNA-menin and AR-RNA-menin co-condensation, plus the fact that AR and MLL have separate binding regions on menin, in principle, enable the formation of condensates incorporating all these components, which might seed and mechanistically explain the buildup of superenhancers targeting AR activity on specific genes [49,50]. Analogous mechanisms can be perceived in other super enhancers, nucleated by Oct4 [51], Sox 2 [52] and Med1 [50,53], regulating other transcription systems [52]. In accord, we have observed extensive cellular colocalization of both menin and MLL in AR foci, forming upon AR overexpression and hormonal activation. A luciferase reporter assay also showed that these events coincide with AR-driven, ARE-specific transcription. In accord, oncogenic events, such as menin upregulation [5,7] or AR-v7 overexpression [54,55], which we have observed to affect dynamics of sub-condensates *in vitro*, may have an effect on the fine tuning of this system *in vivo*.

As oncogenic upregulation of menin - an increase in its concentration - promotes its dimerization, which has different outcomes if AR or MLL is overexpressed in cancer cells, it either inhibits MLL-RNA LLPS or makes AR-RNA condensates much less dynamic (Fig. 10), at least part of the mechanism of these oncogenic events can be the misregulation of the activity and/or specificity of AR condensates. As AR overexpression is characteristic of prostate and breast cancer, whereas MLL overexpression primarily occurs in leukemia (cf. Suppl. Fig. S7, S8), for example, these scenarios may provide cancer-specific insight into disease mechanisms. As cancer-related aberrant condensation events often seem to rely on the upregulation and/or overactivation of transcription driven by condensate formation [50,56,57], here the more likely scenario is the compromise of precise targeting by abrogated phase separation by the aberrant action of menin or AR-v7. Given that highly specific protein-protein interactions occur between both menin-MLL and menin-AR, this creates a new opportunity of interfering with cancer by the rational targeting of these mechanisms [15].

## Abbreviations

AR: androgen receptor; ARE: androgen response element; BLI: bio-layer interferometry; CD: circular dichroism; CRPC: castration-resistant prostate cancer; CS-FBS: charcoal-stripped FBS; CTD: C-terminal domain; DLS: dynamic light scattering; DBD: DNA-binding domain; DHT: dihydrotestosterone; FL: full-length; FRAP: fluorescence recovery after photobleaching; GFP-AR: GFP-fused AR; HPLC: high-performance liquid chromatography; HMT: histone methyltransferase; HSQC: heteronuclear single quantum coherence; IDR: intrinsically disordered region; LBD: ligand-binding domain; LLPS: liquid-liquid phase separation; PDB: Protein data bank; SEC-MALS: size-exclusion chromatography, coupled with multiangle light scattering detector; MBM: menin-binding motif (of MLL); MBR: menin-binding region (of AR); MEN: multiple endocrine neoplasia; MLL: mixed lineage leukemia (protein); MST: microscale thermophoresis; NMR: nuclear magnetic resonance; NTD: N-terminal domain; PC: prostate cancer; PDB: protein data bank; PSA: prostate-specific antigen; SAXS: small-angle X-ray scattering

## Conflict of Interest

The authors declare no conflict of interest.

## Author Contributions

JA formulated concepts and designed experiments carried out biophysical and cellular analyses, bioinformatics calculations and wrote the manuscript. JKC helped with NMR experiments and analysis. AM helped with MST. PT formulated the concept, managed the project, and wrote the manuscript.

## Supporting information

Described in the main text.

## Acknowledgment

This work was supported by grants K124670 and K131702 from the National Research, Development and Innovation Office (NKFIH, Hungary, to PT), a VUB Strategic Research Program on Microfluidics (SRP51) at Vrije Universiteit Brussel (VUB, Brussels, Belgium, to PT), an EC H2020-WIDESPREAD-2020-5 Twinning grant (PhasAge, no. 952334, to PT) and EC H2020-MSCA-RISE Action grant (IDPfun, no. 778247, to PT), and an FWO PhD fellowship in strategic basic research (FWOSB77, to JA).

We are indebted to the help of Dr. Alexandre Wohlkönig for providing purified full-length AR, Dr. Alexander Shkumatov (VIB-VUB Center for Structural Biology, Brussels, Belgium) for his advice on analyzing SAXS results, Jesper Skøttegaard Øemig for discussions on the project and Prof. Alex Volkov with guiding NMR experiments (all from VIB-VUB Center for Structural Biology, Brussels, Belgium).

## Material and methods

### Constructs

Different constructs of menin: Wild-type menin referred to as menin, menin Δ460-519 as menin ΔIDR1 [10], menin Δ460-519/ 586-615 as meninΔ1DR1, IDR2.

AR-v7 was used to generate different constructs: AR-v7 DBD referred to as AR DBD-alt. (aa D552-C649), AR MBR-DBD (aa. G473-C649), AR MBR (aa. G473-T560) [5,12]: AR MBR was used to generate two truncated versions, AR MBR A (aa. G473-T519) and MBR B (aa. R 512-560) have been cloned into pHYRSF53 for expressing SUMO-tagged proteins. AR DBD hinge (aa D 551-T671) was also subcloned to the pHYRSF53 plasmid. Four MLL Peptides of 13, 14, 20, and 29 amino acids, which have been shown to interact with menin, were ordered from Genescipt.

Polyuridylic acid (poly U) was ordered from sigma aldrich. Biotinylated C3(1) gene containing ARE was ordered from eurofin and annealed to measure binding with different constructs of androgen receptor. The sequence of ARE is shown below.

>C3(1)-ARE+flanking [12]

AGCTTACAT**<u>AGTACG</u>**TGA**<u>TGTTCT</u>**CAAGGTCGA

### Protein expression and purification

Purification of MBR, MBR DBD, and DBD have been described before [12], a similar strategy was used for DBD-Hinge and menin. Briefly, these constructs were expressed in BL21-star (DE3) ^TM^ (Thermo Fisher Scientific) with His-sumo tag on the N terminus of the protein. Expression of menin was induced at OD 0.5-0.7 with 0.1 mM isopropyl β-d-1-thiogalactopyranoside (IPTG), at 25 °C shaking overnight while the AR constructs were expressed with 1mM IPTG at 28 °C for four hours, cells were collected, and pellets were resuspended in lysis buffers. Single (^15^N) and double (^15^N, ^13^C) isotopic label proteins were expressed in minimal media with ^15^NH_4_Cl and ^13^C-glucose as Nitrogen and Carbon sources, respectively. M9 media preculture was grown overnight by transferring 1 ml of L.B culture,the next morning this preculture was transferred to M9 media for protein expression.

For menin, lysis buffer (50 mM Tris, 600 mM NaCl, 5 mM benzamidine, 0.5 mM TCEP, pH 8.0, EDTA free protease inhibitors) and for AR (100 mM Tris, 300 mM NaCl, 5% glycerol, 5 mM benzamidine, 0.5 mM TCEP, pH 7.5, EDTA free protease inhibitors). Cells were lysed by sonication (Sonica VCX-70 vibra cell) for 10 minutes (5s pulse on, 5s pulse off, 60% amplification), afterwards DNASE I (5 μg /ml), and cell debris were removed by centrifugation at 40.000X g for an hour. The supernatant was loaded on Histrap Hp ^TM^ 5ml column, proteins were eluted with 500 mM imidazole followed by buffer exchange to buffer (50mM Tris, 300 mM NaCl, pH 8.0) with HiTrap 26/10 Desalting column (GE Healthcare). To obtain protein without sumo tag, the buffer exchanged sample was incubated with ULP1 protease in a cold room for one hour, followed by reversing his trap to obtain His tag-sumo and collect native protein in flow through.

To obtain the His-sumo protein and only His-sumo tag, after buffer exchange the sample was loaded on HiTrap QFF ^TM^ anion exchange column and was eluted with 1M NaCl. Constructs with DBD were purified by using Hitrap ® Heparin High Performance 5 ml column. The last step of purification: Hishtag-sumo-AR, Hitag-sumo were purified by using S75-16/600 (GE healthcare) column equilibrated with 50 mM Tris-base, 150 mM NaCl, 0.5 mM TCEP, pH 8.0 and menin was purified by S200 16/600 (GE healthcare) equilibrated with 50 mM Tris-base, 300 mM NaCl, 0.5 mM TCEP, pH 8.0. Fractions containing protein were collected, concentrated and flash frozen to store at -80 C.

### Phase separation

Different Peptides of MLL (MLL Pep1, MLL Pep 2, MLL Pep 3, MLL Pep 4) and AR constructs were induced to phase separate at 25-50 µM with 0.5 µg/ µl of poly U in buffer (50 mM phosphate, 75 mM NaCl, 0.5 mM TCEP). Proteins and RNA without binding partners were used as a control.

### Turbidity measurements

The turbidity of the 40 μl sample was measured in 384 well black transparent bottom microplates (Greiner bio-one, chimney well μclear ®). After loading and mixing the samples, the plate was covered with a transparent film view seal (Greiner). BioTek SynergyTM Mx plate reader was used to measure absorbance at 600 nm for 12 hr at 25 °C

### Microscopy

Phase separation of MLL and AR in the presence or absence of menin was monitored by Fluorescence microscopy, Labeling was performed by Dylight® 488 dye (Thermo Scientific) and RED-NHS 2nd generation (nano temper) was performed according to manufacturer’s protocol. Labeled proteins were mixed with 200 times excess of unlabelled protein to obtain the final concentration of protein.

Microscopy was performed with Leica DMi8 microscope equipped with a Leica DFC7000 GT camera, 10-20 ul of sample was loaded onto glass slide and covered with cover slip. Droplets were observed by 100X oil immersion objectives.

### Circular Dichroism

Circular dichroism (CD) spectrum of menin, menin ΔIDR1 and AR MBR was acquired in buffer (50 mM phosphate, 10 µM TCEP, pH 7.5). Spectra was obtained by a wavelength scan of 2 μM menin/ menin ΔIDR1 and 7 μM of AR MBR. Data was obtained by Jasco CD Spectro polarimeter at 25 °C by using 1 mm quartz cuvette from Hellma Analytics in the Far UV region (190-260 nm) with continuous scanning mode. Data pitch 0.5 nm, scanning speed 50 nm /min, 2s response time with band width accumulation of 6.

After measurement ellipticity was obtained in milli degree (m deg), these units were converted to mean residue molar ellipticity by using formula from Biozentrum biophysics, the formula is given below.

Θ=(Ellipticity.10^6)/(l.c.n)

Θ Mean residual molar ellipticity [degree cm2.dmol-1] l Path length [cm]

C Protein concentration in [µM] n Number of Peptide bonds [-]

After wavelength scan, thermal scan was carried out at 222 nm wavelength between 25 °C and 95 °C, Ellipticity was plotted against temperature for 2 μM of menin/ menin ΔIDR1. The melting temperature (Tm) was calculated by using formula.

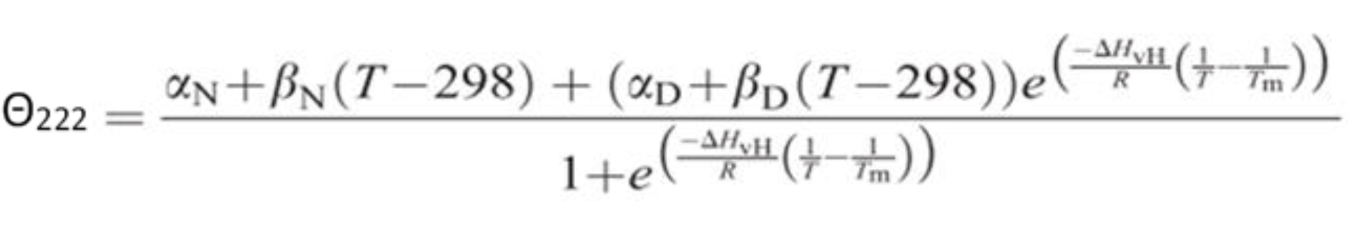

Θ_222_ Observed ellipticity, αN Ellipticity of native states at 25 °C, αD Ellipticity of denatured state at 25°C, β_N_ Slope of native state, β_D_ Slope of denatured states,T Temperature, Tm Midpoint denaturation temperature, R Gas constant, ΔH_vH_ Van’tHofff enthalpy change of unfolding [60].

### Dynamic light scattering

Dynamic light scattering (DLS) measurements were performed on a DynaPro NanoStar (Wyatt) instrument. The sample was centrifuged for 30 minutes at 16,200 X g before measurement. A disposable cuvette (WYATT technology) was filled with 100 ul of menin/ menin ΔIDR1/ menin ΔIDR1, IDR2 and menin-MLL or menin-AR MBR complex with 5-10 X concentration of MLL/ AR MBR. Measurements were carried out at different concentrations of menin to obtain monomer and dimer.The intensity of scattered light was recorded at 25° on 95**°,** collecting 10 acquisitions (8 s each). Measurements were repeated at least three times.The software package used was DYNAMICS 7.1.9 and data was plotted by Graphpad prism.

### Analytical Size-exclusion chromatography

Pre-equilibrated S200 increase 10/300 GL column with buffer (50mM Tris, 150mM NaCl, pH 8.0, 0.5 mM TCEP) was used for analytical gel filtration of menin at different concentrations. The column was loaded with 200 µl of samples. Column calibration was performed by BioRad gel filtration standard having thyroglobulin (bovine) 670 kDa, gamma globulin 158 kDa, oval albumin 44 kDa, myoglobin 17 kDa, vitamin B12 1.3 kDa proteins. By using this standard, the apparent molecular mass of menin was calculated by using

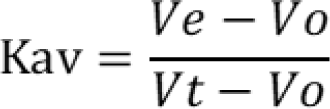

Ve Elution volume of a component Vo Void volume of the column

Vt Total volume of column

Kav Partition coefficient

### Size-exclusion chromatography-Multi Angle Light Scattering

High performance liquid chromatography (HPLC) coupled with a multiangle light scattering detector (SEC-MALS) was used to measure the mass of protein. Agilent Bio SEC - column was pre-equilibrated with a running buffer (50 mM Tris, 150 mM NaCl, 1mM TCEP, pH 8.0) and different concentrations of 30 ul of menin and AR were loaded on the column.The running buffer was filtered two times through a 0.1 μm filter before starting equilibration. Multi angle light scattering setup is composed of an Ultraviolet detector, Shodex RI-501, and TREOS II (Wyatt technology) attached to Agilent Bio SEC -column, molecular mass was calculated by ASTRA 7.1.4 software Wyatt).

### Small-angle X-ray scattering

High performance liquid chromatography (HPLC) coupled to small-angle x-ray scattering (SAXS) and Batch SAXS was collected on SWING (Small and Wide-angle X-ray Scattering). SAXS data was acquired in SOLEIL synchrotron on beamline of SWING (Small and Wide-angle X-ray Scattering) in HPLC and Batch mode at a wavelength of λ=1.03 Å and a AVIEX PCCD170170 detector at distance of 1.8 m, this causes momentum transfer q range of 0.01-0.62 Å^-1^ (λ^−1^; 2θ is scattering angle).

Menin ΔIDR1 in 5 mg/ml and 10 mg/ml (50 ul) was loaded on HPLC Agilent Bio SEC-column equilibrated (50 mM Tris, 300 mM NaCl, 1mM TCEP, pH 8.0) to obtain scattering data. Buffer data was obtained at the start of chromatography and sample data was collected in the peak area. For batch mode, AR MBR (1, 2, 3, 4, 5 mg/ml) was measured with 40 ul volume. Data reduction, Rg evaluation, data averaging and merging was performed with Foxtrot software.ATSAS [61] package was used for data analysis and structure modeling. Scattering curves for both proteins were used for Guinier analysis with Primus Qt [62]. Distance distribution was obtained by GNOM [63].

### Octet Biolayer interferometry

Biolayer interferometry (BLI) was used to study the interaction between AR and menin with and without MLL. Experiments were carried out in a buffer (50 mM Tris, 150 mM NaCl, pH 8.0, 0.5 mM TCEP, 0.05 % tween 20, pH 7.5) on OCTET RED 96. AR MBR /AR MBR A/ AR MBR B was loaded onto Ni-NTA biosensors (FORTBIO) at 5 ug/ml concentration, followed by blocking of sensors with sumo (10 ug /ml), baseline was acquired in buffer followed by association in protein and dissociation in the buffer. The concentration of menin (0.1-50 uM) was used for titration at 25°C with shaking speed of 1000 rpm. In measurement of AR and menin-MLL interaction, after blocking sensors, they were dipped in different concentrations of the menin-MLL complex (0.03-49.7 uM).

Octet signal was obtained firstly in a buffer for 120 sec, followed by loading for 180 sec, then washing for 60 sec before quenching for 120 seconds. The baseline recorded for 120 sec was followed by association and dissociation for 300 and 600 sec, respectively. Each experiment was performed at least three times.The dissociation constant (Kd) was estimated by a fitting response (nm) as a function of protein/protein-protein complex concentration (uM) with Octet data Analysis Software 9.0. The final graph was generated by GraphPad Prism.

### Microscale thermophoresis

The Monolith^TM^ NT.115 instrument (Nano Temper) was used to study homo-dimerisation menin, menin, interaction of menin with AR, and menin-AR-MLL complex. All experiments were performed in a buffer (50 mM Tris, 150 mM NaCl, pH 7.5, 0.05 % Tween 20, 0.5 mM TCEP).

For interaction between menin-AR: menin and AR were labeled with RED-NHS 2nd generation (Nano Temper) by using manufacturing protocol. In the experiment, 50 nM of labeled menin was titrated with an increasing concentration of His-sumo-AR Fragments (MBR, MBR A, MBRB), as negative control His-sumo tag was used. A 2-fold serial dilution of Fragments in the buffer was prepared, with a final volume of 20 ul in each PCR tube. The mixture was loaded to Monolith^TM^NT.115 MST premium coated capillaries. Experiments were carried out at 20 % MST power and 50 % LED power.A similar experiment was also performed with labeled AR MBR.

To determine the dissociation constant of interaction between labeled menin and unlabeled menin, different concentrations of menin were used. In this experiment 50 nM of labeled menin was titrated with increasing concentration of unlabelled menin with 2-fold serial dilution. Then the sample was loaded in premium coated capillaries and measurement was made using Nanotemper with LED power of 20 % and MST power was 50 %.

To determine whether MLL and AR have a different or same binding site on menin, competition experiments were performed, and labeled menin-MLL complex was made in a 1:5 ratio to obtain a complex concentration of 24 nM Two-fold serial dilution of increasing concentration of MBR was titrated with a constant concentration of menin -MLL complex to obtain dissociation constant. The dissociation constant was obtained by using MO. Affinity Analysis software to fit triplicates.

### Nuclear magnetic resonance

NMR spectra were obtained at 600 MHz (Agilent, Coldprobe) and 800 MHz (Bruker Avance III, TCI cryoprobe). Spectra of an AR MBR uniformly labeled with ^15^N were obtained, with a sweep width of 16 ppm in the direct dimension and 35 ppm in the indirect. The number of scans was typically 16. Typical triple-resonance spectra (HNCO, HN(CA)CO, HNCA, HN(CO)CA, HNCACB and CBCA(CO)NH) were utilized for specific backbone resonance assignment at a 1 mM protein concentration. These standard experiments were supplemented with HCACON and HCAN [64] experiments to confirm assignments and to assign the proline residues and those residues where exchange made amide-based experiments unusable. Measurements were performed at 298 k in 50 mM phosphate (pH 6.5), 50mM NaCl, 0.5 mM TCEP and 2.5% D_2_O. Assignments were transferred by titration to 50 mM phosphate (pH 7.0), 300 mM NaCl, 0.5 mM TCEP and 5 %D_2_O buffer.

#### Titration with menin

Titrations were performed by adding concentrated unlabelled menin in the matched buffer to ^15^N-labeled AR iteratively to give concentrations mentioned in the legend.

### Cell culture

Human embryonic kidney (HEK) 293 T, U20S cell lines were maintained in Dulbecco’ modified Eagle’s medium (DMEM) from GIBCO supplemented with Fetal bovine serum (FBS) from GIBCO at 10% and 1% penicillin/ streptavidin from GIBCO. LNCAP cells from ATCC were grown in Rpmi 1640 Medium media. Cells were transfected with Lipofectamine 3000 from Thermo scientific. All cells were maintained at 37 C at 10% CO2 in humidified incubator.

### Antibodies

Primary mouse monoclonal anti-menin (Cat #MA5-38572) antibody purchased from Thermo Fisher Scientific and primary Anti MLL produced in rabbit (Cat # HPA044910) from Merck. Fluorescein (FITC)–conjugated Affinipure Goat Anti-Rabbit IgG(H+L) (Cat #S A00003-2) from proteintech, Cy5 conjugated Goat anti-mouse IgG (H+L) Secondary Antibody (Cat # A10524) and Cy5 conjugated Goat anti-Rabbit IgG (H+L) Secondary Antibody (Cat # A10523) were purchased from Invitrogen.

### Luciferase reporter assay

To explore the effect of AR-dependent transcriptional activity resulting from the formation of condensates. HEK293T cells were transfected with Lipofectamine 3000 reagent (Invitrogen) in Rpmi 1640 Medium media from Thermo scientific with different amounts of pARE-E1b-luc, plasmid was gift from Carlo Rinaldi [65] along with AR FL plasmid. After 24 hr transfection cells were washed with PBS and treated with 10 nM DHT or an equal amount of ethanol. Then after another 24 hr cells were lysed and luciferase activity was measured by using the luciferase assay system (Cat no E4030) from Promega.

### Immunofluorescence

For immunostaining, cells were grown in Ibidi USA µ-Slide VI 0.4, ibiTreat (Cat no 50-305-784) according to manufacturer protocol and starved in 10% charcoal-stripped FBS (CS-FBS) for two days, followed by 10 nM DHT treatment for 24 hr. Cells were fixed with 4% paraformaldehyde (PFA) at room temperature for 15-20 minutes, followed by incubation with respective primary antibody for 1 hr at room temperature followed by fluorescent conjugated secondary antibody and incubate for 1 hr at room temperature, followed by NucBlue™ Fixed Cell ReadyProbes™ Reagent (DAPI) from Thermo Fisher Scientific 2 drops/ml for 15 to 30 minutes. Imagining was done by RCM 2 confocal microscope.

### Alpha Fold predictions

Alphafold2 using MMseqs2 was used for the prediction of the dimeric menin with uniprotKB entry o00255. (https://colab.research.google.com/github/sokrypton/ColabFold/blob/main/AlphaFold2.ipynb).

## Notes

### Competing Interest Statement

The authors have declared no competing interest.

## References

1. Gregory CW, He B, Johnson RT, Ford OH, Mohler JL, French FS, et al. A mechanism for androgen receptor-mediated prostate cancer recurrence after androgen deprivation therapy. Cancer Res. 2001;61: 4315–4319.

2. Papaioannou M, Reeb C, Asim M, Dotzlaw H, Baniahmad A. Co-activator and co-repressor interplay on the human androgen receptor. Andrologia. 2005;37: 211–212.

3. Kim S, Au CC, Jamalruddin MAB, Abou-Ghali NE, Mukhtar E, Portella L, et al. AR-V7 exhibits non-canonical mechanisms of nuclear import and chromatin engagement in castrate-resistant prostate cancer. Elife. 2022;11. doi:10.7554/eLife.73396

4. Cierpicki T, Grembecka J. Challenges and opportunities in targeting the menin–MLL interaction. Future Med Chem. 2014;6: 447–462.

5. Malik R, Khan AP, Asangani IA, Cieślik M, Prensner JR, Wang X, et al. Targeting the MLL complex in castration-resistant prostate cancer. Nat Med. 2015;21: 344–352.

6. Issa GC, Ravandi F, DiNardo CD, Jabbour E, Kantarjian HM, Andreeff M. Therapeutic implications of menin inhibition in acute leukemias. Leukemia. 2021;35: 2482–2495.

7. Cherif C, Nguyen DT, Paris C, Le TK, Sefiane T, Carbuccia N, et al. Menin inhibition suppresses castration-resistant prostate cancer and enhances chemosensitivity. Oncogene. 2021. doi:10.1038/s41388-021-02039-2

8. Park SH, Ayoub A, Lee Y-T, Xu J, Kim H, Zheng W, et al. Cryo-EM structure of the human MLL1 core complex bound to the nucleosome. Nat Commun. 2019;10: 5540.

9. Grembecka J, Belcher AM, Hartley T, Cierpicki T. Molecular basis of the mixed lineage leukemia-menin interaction: implications for targeting mixed lineage leukemias. J Biol Chem. 2010;285: 40690–40698.

10. Huang J, Gurung B, Wan B, Matkar S, Veniaminova NA, Wan K, et al. The same pocket in menin binds both MLL and JUND but has opposite effects on transcription. Nature. 2012;482: 542–546.

11. Lei H, Zhang S-Q, Fan S, Bai H-R, Zhao H-Y, Mao S, et al. Recent Progress of Small Molecule Menin–MLL Interaction Inhibitors as Therapeutic Agents for Acute Leukemia. J Med Chem. 2021;64: 15519–15533.

12. Ahmed J, Meszaros A, Lazar T, Tompa P. DNA-binding domain as the minimal region driving RNA-dependent liquid-liquid phase separation of androgen receptor. Protein Sci. 2021;30: 1380–1392.

13. Zhang F, Wong S, Lee J, Lingadahalli S, Wells C, Saxena N, et al. Dynamic phase separation of the androgen receptor and its coactivators to regulate gene expression. bioRxiv. 2021. p. 2021.03.27.437301. doi:10.1101/2021.03.27.437301

14. Rizo J, Roggero CM, Esser V, Duan L, Rice AM, Ma S. Poly-glutamine-dependent self-association as a potential mechanism for regulation of androgen receptor activity. bioRxiv. 2021. Available: https://www.biorxiv.org/content/10.1101/2021.10.08.463684.abstract

15. Mitrea DM, Mittasch M, Gomes BF, Klein IA, Murcko MA. Modulating biomolecular condensates: a novel approach to drug discovery. Nat Rev Drug Discov. 2022;21: 841– 862.

16. Mészáros B, Erdos G, Dosztányi Z. IUPred2A: context-dependent prediction of protein disorder as a function of redox state and protein binding. Nucleic Acids Res. 2018;46: W329–W337.

17. Erdős G, Dosztányi Z. Analyzing Protein Disorder with IUPred2A. Curr Protoc Bioinformatics. 2020;70: e99.

18. Reid J, Kelly SM, Watt K, Price NC, McEwan IJ. Conformational analysis of the androgen receptor amino-terminal domain involved in transactivation: influence of structure-stabilizing solutes and protein-protein interactions. J Biol Chem. 2002;277: 20079–20086.

19. Shaffer PL, Jivan A, Dollins DE, Claessens F, Gewirth DT. Structural basis of androgen receptor binding to selective androgen response elements. Proc Natl Acad Sci U S A. 2004;101: 4758–4763.

20. Nadal M, Prekovic S, Gallastegui N, Helsen C, Abella M, Zielinska K, et al. Structure of the homodimeric androgen receptor ligand-binding domain. Nat Commun. 2017;8: 14388.

21. Wasmuth EV, Broeck AV, LaClair JR, Hoover EA, Lawrence KE, Paknejad N, et al. Allosteric interactions prime androgen receptor dimerization and activation. Mol Cell. 2022;82: 2021–2031.e5.

22. Peacock SO, Fahrenholtz CD, Burnstein KL. Vav3 enhances androgen receptor splice variant activity and is critical for castration-resistant prostate cancer growth and survival. Mol Endocrinol. 2012;26: 1967–1979.

23. Magani F, Peacock SO, Rice MA, Martinez MJ, Greene AM, Magani PS, et al. Targeting AR Variant–Coactivator Interactions to Exploit Prostate Cancer Vulnerabilities. Molecular Cancer Research. 2017. pp. 1469–1480. doi:10.1158/1541-7786.mcr-17-0280

24. Gray FLV, Murai MJ, Grembecka J, Cierpicki T. Detection of disordered regions in globular proteins using ^13^C-detected NMR. Protein Sci. 2012;21: 1954–1960.

25. Tompa P. Intrinsically unstructured proteins. Trends Biochem Sci. 2002;27: 527–533.

26. Shkumatov AV, Chinnathambi S, Mandelkow E, Svergun DI. Structural memory of natively unfolded tau protein detected by small-angle X-ray scattering. Proteins: Structure, Function, and Bioinformatics. 2011. pp. 2122–2131. doi:10.1002/prot.23033

27. Camilloni C, De Simone A, Vranken WF, Vendruscolo M. Determination of secondary structure populations in disordered states of proteins using nuclear magnetic resonance chemical shifts. Biochemistry. 2012;51: 2224–2231.

28. Micsonai A, Wien F, Bulyáki É, Kun J, Moussong É, Lee Y-H, et al. BeStSel: a web server for accurate protein secondary structure prediction and fold recognition from the circular dichroism spectra. Nucleic Acids Res. 2018;46: W315–W322.

29. Jarzab A, Kurzawa N, Hopf T, Moerch M, Zecha J, Leijten N, et al. Meltome atlas—thermal proteome stability across the tree of life. Nat Methods. 2020;17: 495–503.

30. Jumper J, Evans R, Pritzel A, Green T, Figurnov M, Ronneberger O, et al. Highly accurate protein structure prediction with AlphaFold. Nature. 2021;596: 583–589.

31. Krissinel E, Henrick K. Inference of macromolecular assemblies from crystalline state. J Mol Biol. 2007;372: 774–797.

32. Jiang H. The complex activities of the SET1/MLL complex core subunits in development and disease. Biochim Biophys Acta Gene Regul Mech. 2020;1863: 194560.

33. Kumar M, Gromiha MM, Raghava GPS. Prediction of RNA binding sites in a protein using SVM and PSSM profile. Proteins. 2008;71: 189–194.

34. Eftekharzadeh B, Banduseela VC, Chiesa G, Martínez-Cristóbal P, Rauch JN, Nath SR, et al. Hsp70 and Hsp40 inhibit an inter-domain interaction necessary for transcriptional activity in the androgen receptor. Nat Commun. 2019;10: 3562.

35. Xie J, He H, Kong W, Li Z, Gao Z, Xie D, et al. Targeting androgen receptor phase separation to overcome antiandrogen resistance. Nat Chem Biol. 2022. doi:10.1038/s41589-022-01151-y

36. Lin Y, Mori E, Kato M, Xiang S, Wu L, Kwon I, et al. Toxic PR Poly-Dipeptides Encoded by the C9orf72 Repeat Expansion Target LC Domain Polymers. Cell. 2016;167: 789–802.e12.

37. Perdikari TM, Murthy AC, Fawzi NL. Molecular insights into the effect of alkanediols on FUS liquid-liquid phase separation. bioRxiv. 2022. p. 2022.05.05.490812. doi:10.1101/2022.05.05.490812

38. Olausson, Nistér, Lindström. Loss of nucleolar histone chaperone NPM1 triggers rearrangement of heterochromatin and synergizes with a deficiency in DNA methyltransferase DNMT3A to …. J Biol Chem. Available: https://www.jbc.org/article/S0021-9258(19)56118-X/abstract

39. Siemund AL, Hanewald T, Kowarz E, Marschalek R. MLL-AF4 and a murinized pSer-variant thereof are turning on the nucleolar stress pathway. Cell Biosci. 2022;12: 47.

40. Vélot L, Lessard F, Bérubé-Simard F-A, Tav C, Neveu B, Teyssier V, et al. Proximity-dependent Mapping of the Androgen Receptor Identifies Kruppel-like Factor 4 as a Functional Partner. Mol Cell Proteomics. 2021;20. doi:10.1016/j.mcpro.2021.100064

41. Russo JW, Nouri M, Balk SP. Androgen Receptor Interaction with Mediator Complex Is Enhanced in Castration-Resistant Prostate Cancer by CDK7 Phosphorylation of MED1. Cancer discovery. 2019. pp. 1490–1492.

42. van de Wijngaart DJ, Dubbink HJ, van Royen ME, Trapman J, Jenster G. Androgen receptor coregulators: recruitment via the coactivator binding groove. Mol Cell Endocrinol. 2012;352: 57–69.

43. Kim T, Jeong K, Kim E, Yoon K, Choi J, Park JH, et al. Menin Enhances Androgen Receptor-Independent Proliferation and Migration of Prostate Cancer Cells. Mol Cells. 2022;45: 202–215.

44. Motlagh HN, Wrabl JO, Li J, Hilser VJ. The ensemble nature of allostery. Nature. 2014;508: 331–339.

45. Hilser VJ. Biochemistry. An ensemble view of allostery. Science. 2010. pp. 653–654.

46. Tompa P. Multisteric regulation by structural disorder in modular signaling proteins: an extension of the concept of allostery. Chem Rev. 2014;114: 6715–6732.

47. Hilser VJ, Thompson EB. Structural dynamics, intrinsic disorder, and allostery in nuclear receptors as transcription factors. J Biol Chem. 2011;286: 39675–39682.

48. Eteleeb AM, Thunuguntla PK, Gelev KZ, Tang CY, Rozycki EB, Miller A, et al. LINC00355 regulates p27KIP expression by binding to MENIN to induce proliferation in late-stage relapse breast cancer. NPJ Breast Cancer. 2022;8: 49.

49. Chen X, Ma Q, Shang Z, Niu Y. Super-enhancer in prostate cancer: transcriptional disorders and therapeutic targets. NPJ Precis Oncol. 2020;4: 31.

50. Sabari BR, Dall’Agnese A, Boija A, Klein IA, Coffey EL, Shrinivas K, et al. Coactivator condensation at super-enhancers links phase separation and gene control. Science. 2018;361. doi:10.1126/science.aar3958

51. Takayama K-I, Kosaka T, Suzuki T, Hongo H, Oya M, Fujimura T, et al. Subtype-specific collaborative transcription factor networks are promoted by OCT4 in the progression of prostate cancer. Nat Commun. 2021;12: 3766.

52. Hnisz D, Abraham BJ, Lee TI, Lau A, Saint-André V, Sigova AA, et al. Super-enhancers in the control of cell identity and disease. Cell. 2013;155: 934–947.

53. Whyte WA, Orlando DA, Hnisz D, Abraham BJ, Lin CY, Kagey MH, et al. Master transcription factors and mediator establish super-enhancers at key cell identity genes. Cell. 2013;153: 307–319.

54. Chen X, Bernemann C, Tolkach Y, Heller M, Nientiedt C, Falkenstein M, et al. Overexpression of nuclear AR-V7 protein in primary prostate cancer is an independent negative prognostic marker in men with high-risk disease receiving adjuvant therapy. Urologic Oncology: Seminars and Original Investigations. 2018;36: 161.e19–161.e30.

55. Zheng Z, Li J, Liu Y, Shi Z, Xuan Z, Yang K, et al. The Crucial Role of AR-V7 in Enzalutamide-Resistance of Castration-Resistant Prostate Cancer. Cancers . 2022;14. doi:10.3390/cancers14194877

56. Cho W-K, Spille J-H, Hecht M, Lee C, Li C, Grube V, et al. Mediator and RNA polymerase II clusters associate in transcription-dependent condensates. Science. 2018;361: 412–415.

57. Boija A, Klein IA, Sabari BR, Dall’Agnese A, Coffey EL, Zamudio AV, et al. Transcription Factors Activate Genes through the Phase-Separation Capacity of Their Activation Domains. Cell. 2018;175: 1842–1855.e16.

58. Uhlén M, Fagerberg L, Hallström BM, Lindskog C, Oksvold P, Mardinoglu A, et al. Proteomics. Tissue-based map of the human proteome. Science. 2015;347: 1260419.

59. Wang M, Weiss M, Simonovic M, Haertinger G, Schrimpf SP, Hengartner MO, et al. PaxDb, a database of protein abundance averages across all three domains of life. Mol Cell Proteomics. 2012;11: 492–500.

60. Sehgal P, Otzen DE. Thermodynamics of unfolding of an integral membrane protein in mixed micelles. Protein Sci. 2006;15: 890–899.

61. Franke D, Petoukhov MV, Konarev PV, Panjkovich A, Tuukkanen A, Mertens HDT, et al. ATSAS 2.8: a comprehensive data analysis suite for small-angle scattering from macromolecular solutions. J Appl Crystallogr. 2017;50: 1212–1225.

62. Konarev PV, Volkov VV, Sokolova AV, Koch MHJ, Svergun DI. PRIMUS: a Windows PC-based system for small-angle scattering data analysis. J Appl Crystallogr. 2003;36: 1277– 1282.

63. Svergun DI. Determination of the regularization parameter in indirect-transform methods using perceptual criteria. J Appl Crystallogr. 1992;25: 495–503.

64. Kanelis V, Donaldson L, Muhandiram DR, Rotin D, Forman-Kay JD, Kay LE. Sequential assignment of proline-rich regions in proteins: application to modular binding domain complexes. J Biomol NMR. 2000;16: 253–259.

65. Lim WF, Forouhan M, Roberts TC, Dabney J, Ellerington R, Speciale AA, et al. Gene therapy with AR isoform 2 rescues spinal and bulbar muscular atrophy phenotype by modulating AR transcriptional activity. Sci Adv. 2021;7. doi:10.1126/sciadv.abi6896

