## Supplementary material for "Menin regulates androgen receptor- and MLL-driven condensation, upsetting regulation of cellular AR-driven transcription": Described in the main text.: Supplementry Figures.pdf

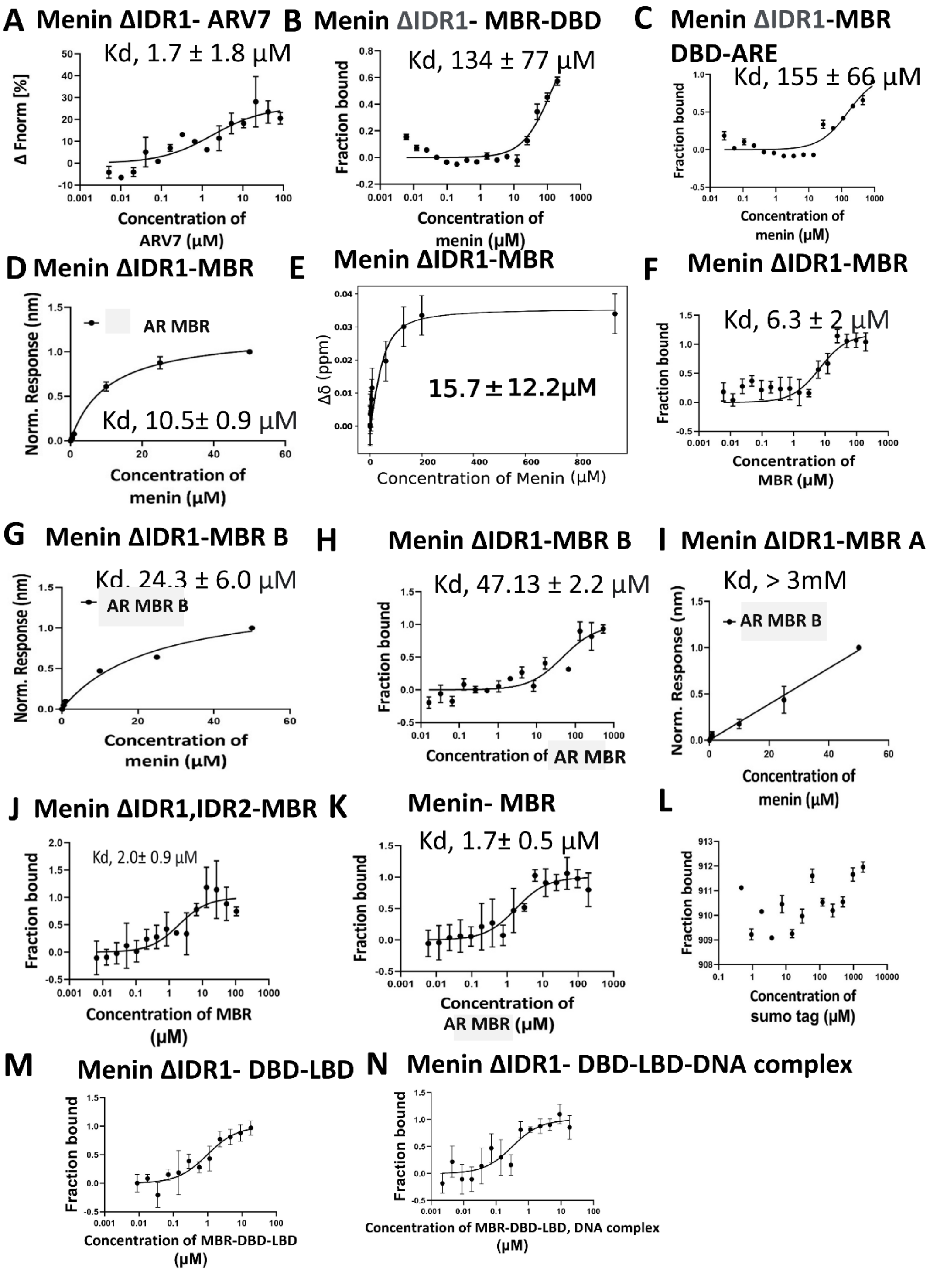


**Suppl. Fig. S1** *Interactions within the menin-AR-MLL complex*

Menin and menin mutant interaction with A) AR-v7 B) MBR-DBD C) MBR-DBD-ARE D) MBR E) MBR F) MBR G)MBR-B H) MBR-B I) MBR-A J) MBR K) MBR L) Sumo tag M) MBR-DBD-LBD, N)MBR-DBD-LBD -DNA complex by MST, BLI and NMR.

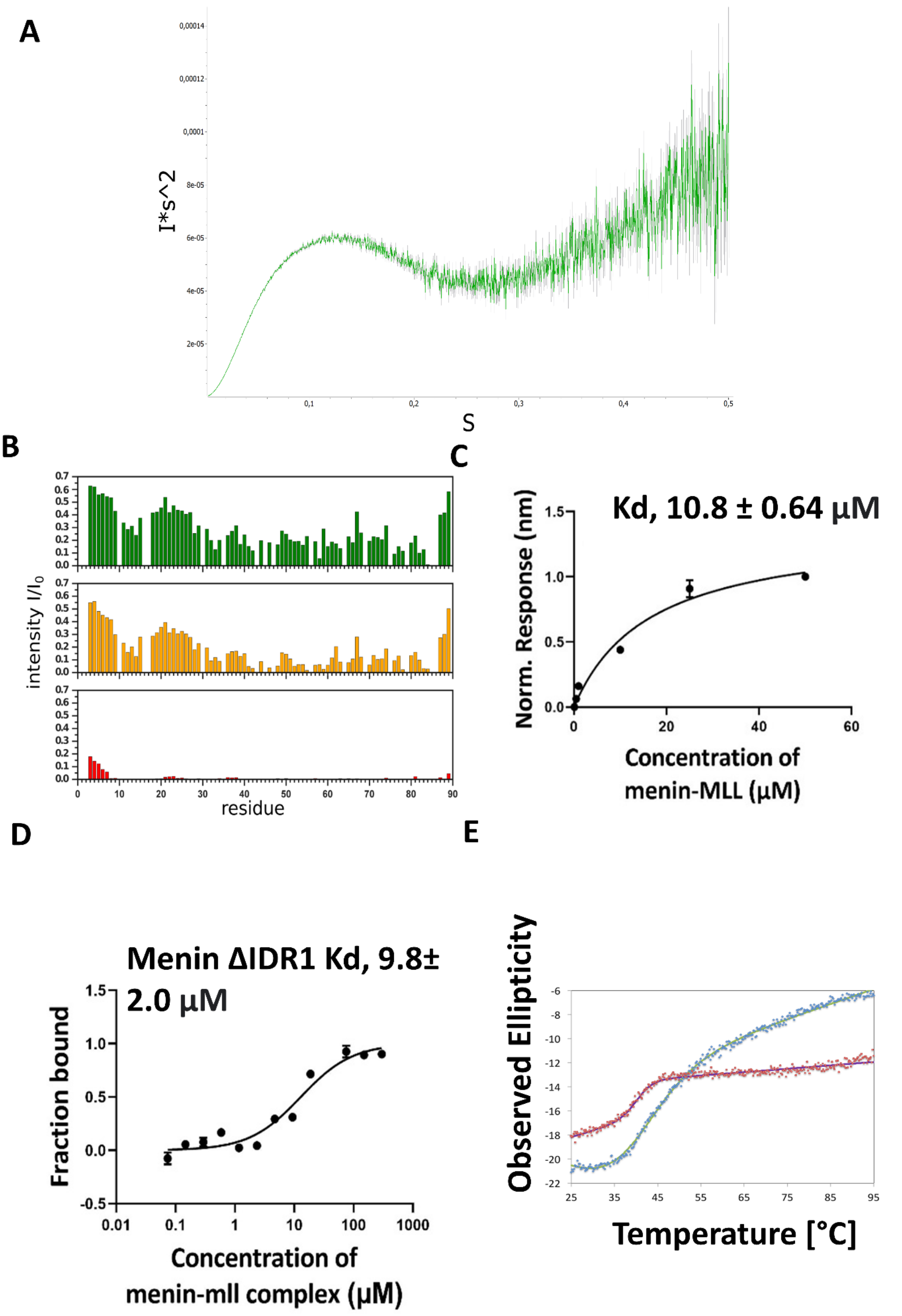


**Suppl. Fig. S2** *menin, AR and menin-AR interaction characterisation*

A) Kratky plot for AR MBR obtained at concentrations 5 mg/ml, B) Decrease in intensity of AR MBR upon addition of menin at different concentrations,C) Interaction of AR with menin- MLL complex by BLI, D) MST E) Thermal scan at 222 nm by CD for menin (green), menin ΔIDR1(red) at 2 μM concentration

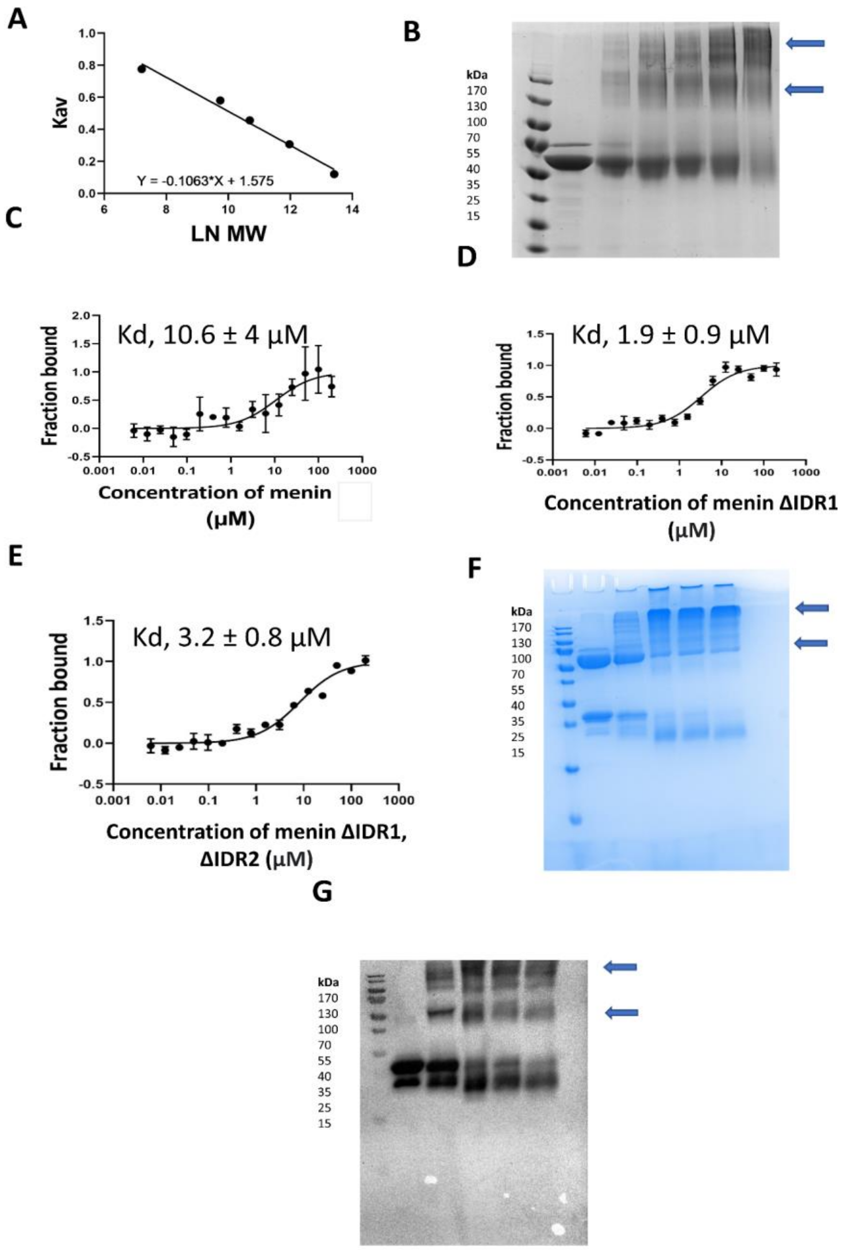


**Suppl. Fig. S3** *menin-AR interaction by MST and crosslinking*

(A) Calibration plot of the natural logarithm of molar mass (Ln MW) and partition coefficient to calculate apparent molar mass of menin. (B) 10 % SDS-PAGE gel for cross-linking of menin at 15 μM in the absence (Lane 1) and presence of a different concentration of glutaraldehyde (0.01, 0.03, 0.05,0.1,0.5) for lane 2 to 6.Interaction of different constructs menin with (C) menin (D) menin ΔIDR1 (E) menin ΔIDR1, IDR2 (F). SDS-PAGE for menin at 15 μM in presence of 15 μM AR MBR, cross-linking by glutaraldehyde stopped after 15, 30, 45, and 60 minutes for lanes 2 to 6, sample but for lane 1 was obtained at time zero G) western blot for F)

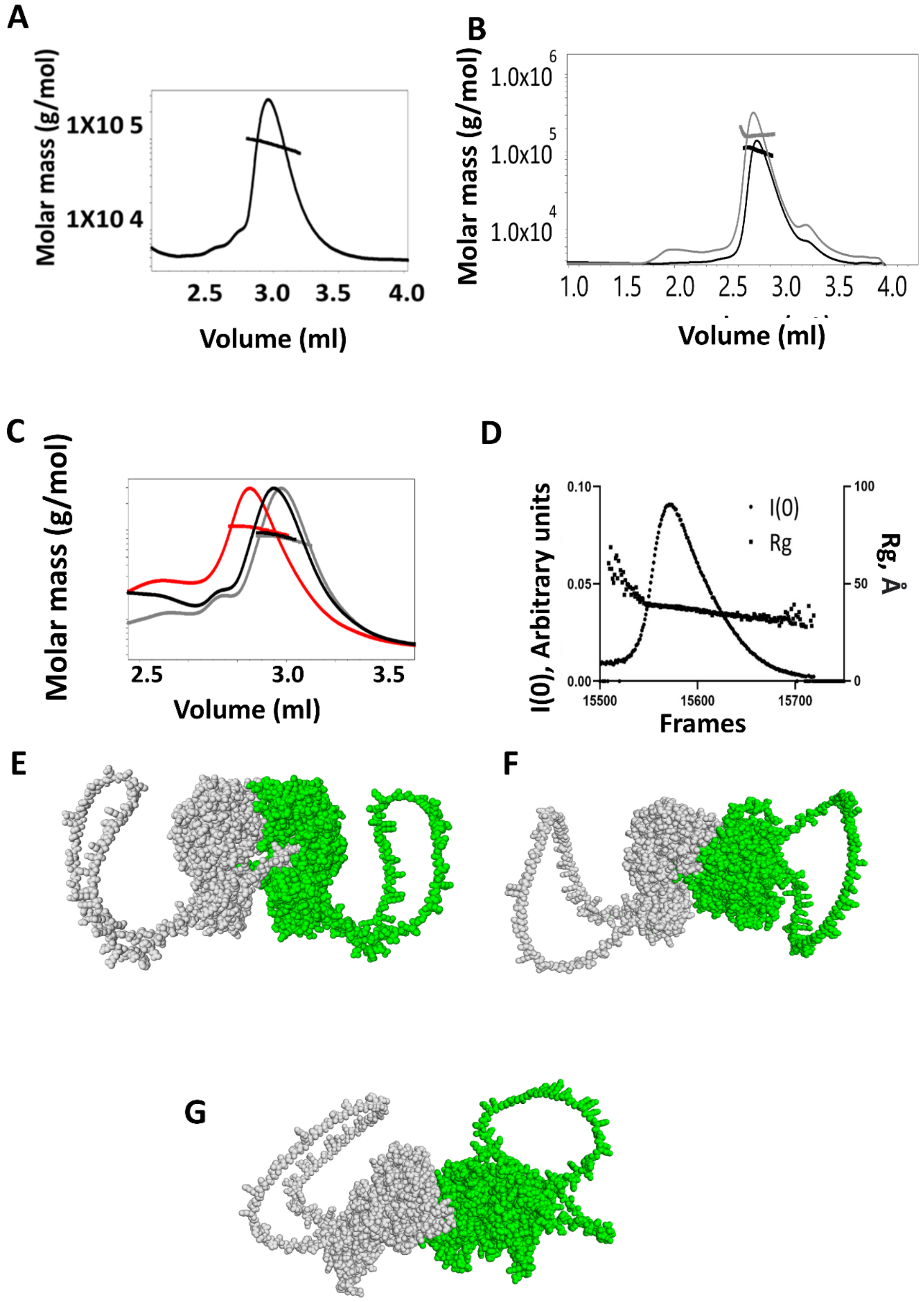


**Suppl. Fig. S4 Menin** *dimeric state and alpha fold predictions for dimer*

(A-C) SEC-MALS of menin and menin ΔIDR1 by using Agilent Bio SEC - column. (H) menin ΔIDR1 at 5 mg/ml, (I) menin at 10 mg/ml (black), and 30 mg/ml (gray), (J) menin ΔIDR1,IDR2 at 2.5 mg/ml (gray), 10 mg/ml (black), and 30 mg/ml (red).
D) SEC-SAXS of menin by using Agilent Bio SEC column at 5 mg/ml to obtain Rg and molar mass of menin .
E), F), G) Different dimeric models predicted by using alpha fold for menin.

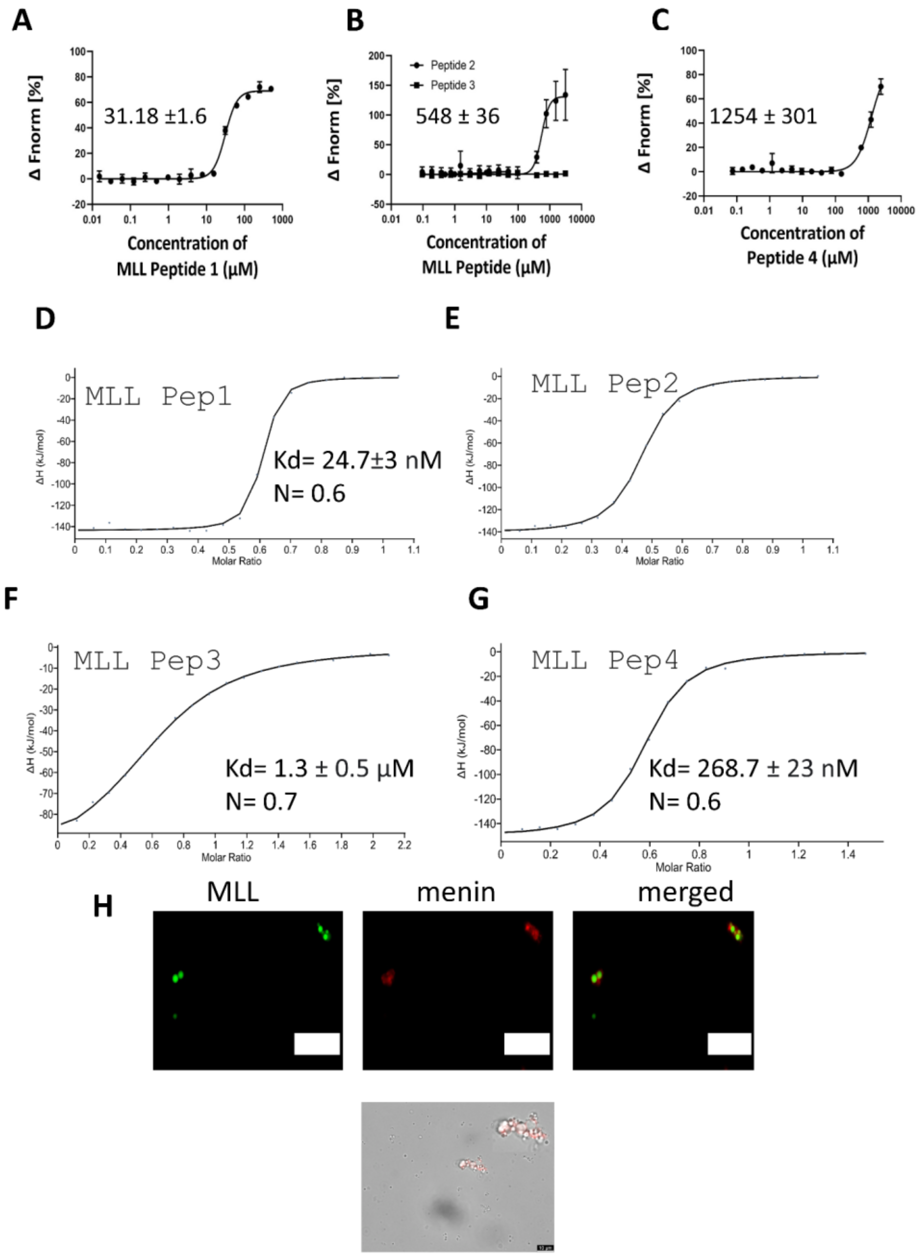


**Suppl. Fig. S5** *Menin interaction with MLL and RNA*
Interaction of different MLL Peptides with RNA A, B, C) and menin D,E,F,G) by microscale thermophoresis (MST) and isothermal titration calorimetry (ITC), OD 600 for Peptide 4 at different concentrations 0 to 100 µM. F) Microscopic image of MLL droplets in presence of menin.

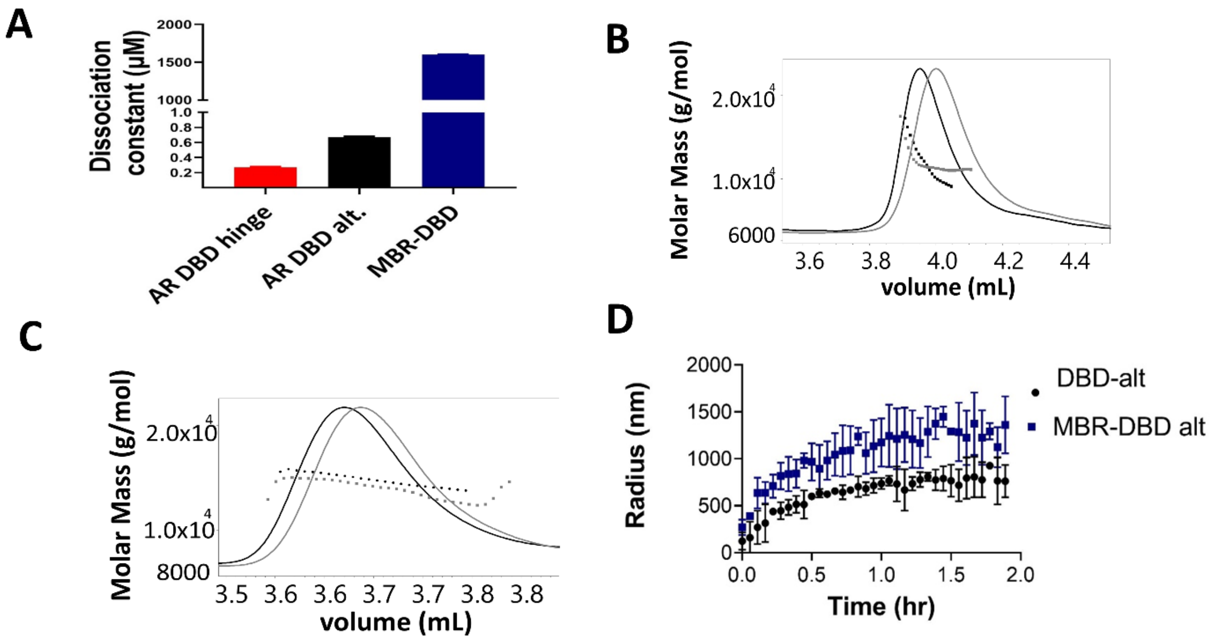


**Suppl. Fig. S6** *AR interaction with ARE DNA and phase separation of AR*

A) Interaction of ARE DNA with AR constructs, measured by BLI

(B, C) SEC-MALS of AR DBD-alt hinge (B), and DBD-hinge (C)

D) Dynamic light scattering (DLS) for 25 µM of DBD and MBR DBD in presence of RNA at 0.05 µg/ µl. Increase in droplet size as function of time is plotted

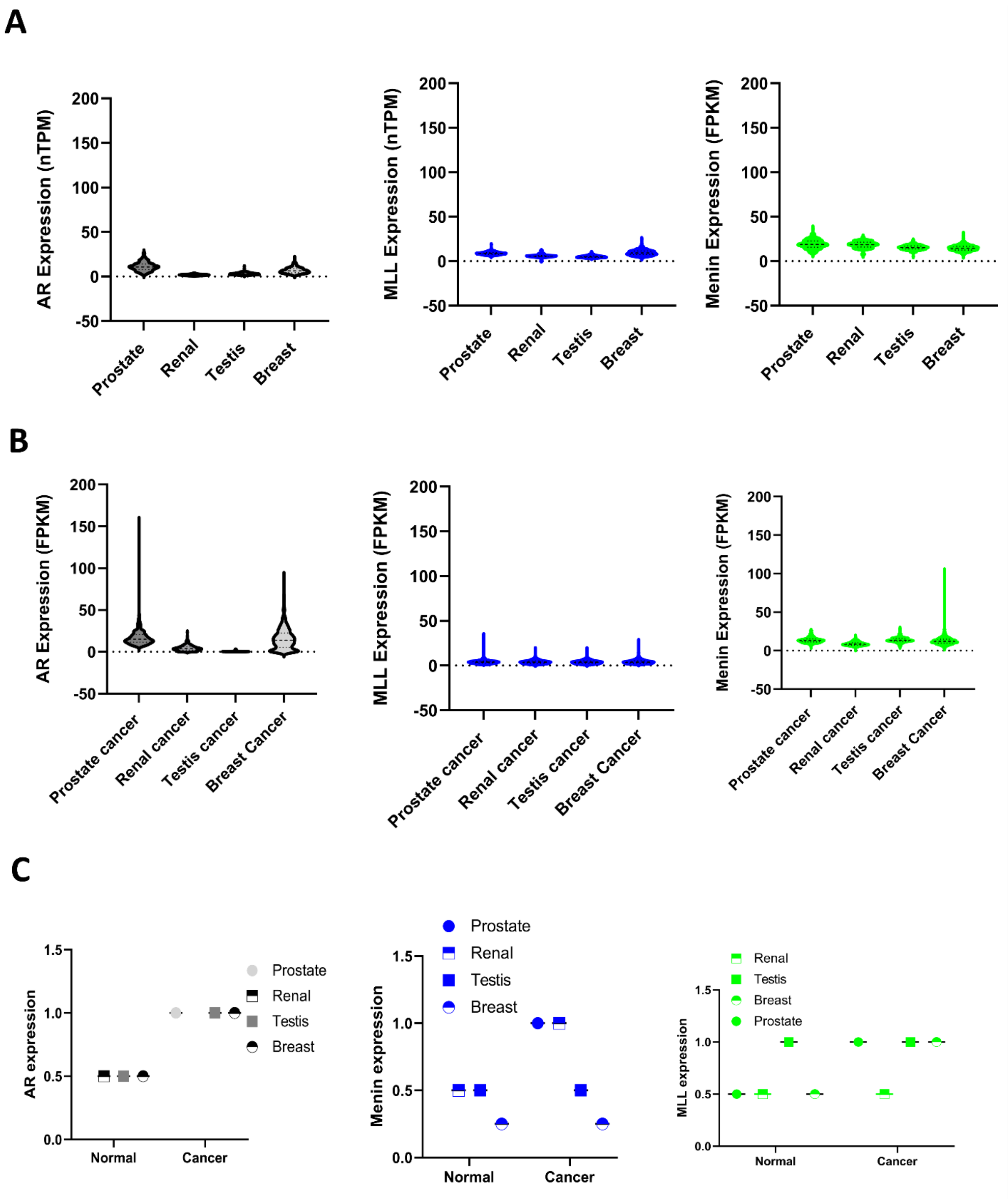


**Suppl. Fig. S7** *AR interaction in different cancers*

*A) Transcript level expression of AR menin and MLL in normal Prostate, Renal, Testis and Breast B) Cancer*

*C) Protein level expression of menin, AR and MLL in different normal vs cancerous tissues.*

*Data was obtained from [58]*

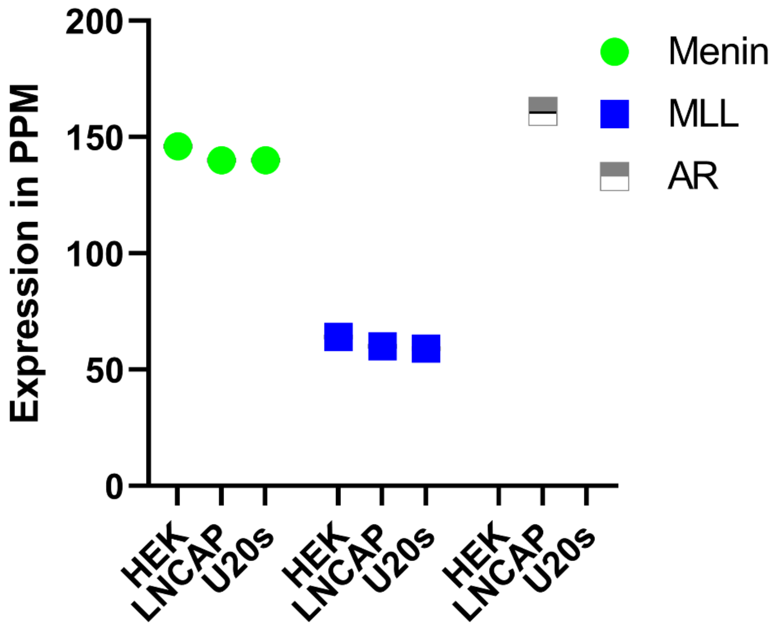


**Suppl. Fig. S8** *AR menin and MLL expression in different cell lines obtained from Protein Abundance Database(paxdb)[59].*

**Suppl. Table S1** *Chemical shifts of AR MBR*

| | | H | NH | C | Cα | Cβ | | |
|---|---|---|---|---|---|---|---|---|
| 2 | Gly | 8.40 | 109.76 | 174.33 | 45.03 | - | 3.74:Hαa | |
| 3 | Gly | 8.21 | 109.14 | 174.09 | 44.83 | - | 3.62:Hαa | |
| 4 | Glu | - | - | 176.40 | - | - | | |
| 4 | Glu | 8.20 | 120.75 | - | 56.30 | 29.69 | 3.92:Hα | |
| 5 | Ala | 8.26 | 125.30 | 178.11 | 52.54 | 18.58 | 3.95:Hα | |
| 6 | Gly | 8.11 | 108.14 | 173.47 | 44.74 | - | 3.58:Hαa | |
| 7 | Ala | 7.78 | 123.54 | 177.44 | 52.02 | 19.02 | 3.99:Hα | |
| 8 | Val | 7.90 | 119.82 | 175.36 | 61.57 | 32.55 | 3.73:Hα | |
| 9 | Ala | 8.15 | 129.51 | 175.14 | 50.02 | 17.76 | 4.15:Hα | |
| 10 | Pro | - | 134.82 | - | 62.75 | 31.55 | 4.05:Hα | |
| 11 | Tyr | 7.98 | 120.13 | 176.30 | 57.99 | - | 4.11:Hα | |
| 12 | Gly | 7.97 | 111.37 | 173.21 | 44.75 | - | 3.37:Hαa | 3.48:Hαb |
| 13 | Tyr | 7.63 | 120.14 | 175.40 | 57.73 | 38.73 | 4.22:Hα | |
| 14 | Thr | 7.75 | 118.09 | 172.97 | 61.09 | 69.79 | 3.90:Hα | |
| 15 | Arg | 8.06 | 125.30 | 173.35 | 53.70 | 29.84 | 4.13:Hα | |
| 16 | Pro | - | 138.54 | - | 61.14 | - | 4.28:Hα | |
| 17 | Pro | - | 135.44 | 176.58 | 62.77 | 31.63 | 4.04:Hα | |
| 18 | Gln | 8.26 | 120.25 | 176.18 | 55.56 | 29.44 | 3.96:Hα | |
| 19 | Gly | 8.18 | 110.04 | - | - | - | | |
| 19 | Gly | - | - | 173.69 | 44.87 | - | 3.58:Hαa | 3.60:Hαb |
| 20 | Leu | 7.91 | 121.59 | 177.07 | 54.67 | 42.13 | 4.00:Hα | |
| 21 | Ala | 8.13 | 124.62 | 178.07 | 52.46 | 18.62 | 3.96:Hα | |
| 22 | Gly | 8.21 | 108.63 | - | - | - | | |
| 22 | Gly | - | - | 173.99 | 45.01 | - | 3.58:Hαa | 3.63:Hαb |
| 23 | Gln | 7.92 | - | 175.93 | - | - | | |
| 23 | Gln | - | 119.38 | - | 55.28 | 29.23 | 4.03:Hα | |
| 24 | Glu | 8.42 | 122.05 | - | 56.79 | 29.60 | 3.92:Hα | |
| 24 | Glu | - | - | 176.49 | - | - | | |
| 25 | Ser | 8.07 | 115.92 | - | - | - | | |
| 25 | Ser | - | - | 173.75 | 58.16 | 63.55 | 4.05:Hα | |
| 26 | Asp | 8.01 | 121.85 | 175.59 | 53.96 | 40.59 | 4.23:Hα | |
| 27 | Phe | - | - | 175.36 | 57.57 | 39.17 | 4.29:Hα | |
| 27 | Phe | 7.87 | 120.49 | - | - | - | | |
| 28 | Thr | 7.75 | 116.96 | 173.10 | 61.33 | 69.69 | 3.88:Hα | |
| 29 | Ala | 8.02 | 128.28 | 175.16 | 50.23 | 17.83 | 4.13:Hα | |
| 30 | Pro | - | 134.87 | 176.22 | 62.70 | 31.73 | 4.01:Hα | |
| 31 | Asp | 8.11 | 119.74 | 175.75 | 54.09 | 40.64 | | |
| 32 | Val | 7.62 | 119.48 | 174.84 | 61.83 | 32.70 | 3.71:Hα | |
| 33 | Trp | 7.92 | 124.77 | 174.65 | 57.04 | 29.56 | 4.23:Hα | |
| 34 | Tyr | 7.46 | 123.44 | - | 54.90 | 38.34 | 4.27:Hα | |
| 35 | Pro | - | 136.29 | 177.29 | 63.23 | 31.41 | 3.77:Hα | |
| 36 | Gly | 7.95 | 109.18 | 174.50 | 45.02 | - | 3.53:Hαa | |
| 37 | Gly | 7.86 | 108.35 | 173.62 | 44.82 | - | | |
| 38 | Met | 7.94 | 119.55 | 175.91 | 55.17 | - | | |
| 38 | Met | - | - | - | - | 32.55 | 4.08:Hα | |
| 39 | Val | 7.90 | 121.34 | 175.73 | 61.98 | 32.49 | | |
| 40 | Ser | 8.07 | 119.42 | 173.98 | 57.81 | 63.46 | 4.09:Hα | |
| 41 | Arg | - | - | - | - | 30.45 | | |
| 41 | Arg | 8.14 | 123.38 | 175.71 | 55.61 | - | 3.99:Hα | |
| 42 | Val | 7.85 | - | 174.10 | - | 32.13 | | |
| 42 | Val | - | 122.34 | - | 59.47 | - | 4.00:Hα | |
| 43 | Pro | - | 138.40 | - | 62.73 | 31.69 | 3.98:Hα | |
| 44 | Tyr | - | - | - | 55.39 | - | 4.40:Hα | |
| 44 | Tyr | 7.82 | 121.25 | - | - | 37.82 | | |
| 45 | Pro | - | 137.24 | 176.10 | 62.53 | 31.56 | 4.06:Hα | |
| 46 | Ser | 8.09 | 117.64 | 176.69 | 56.10 | 63.02 | 4.40:Hα | |
| 47 | Pro | - | 137.70 | - | 63.20 | 31.70 | 4.15:Hα | |
| 48 | Thr | 7.92 | 113.31 | - | 61.55 | 69.33 | | |
| 49 | Cys | 7.97 | 121.56 | - | 58.05 | 27.79 | 4.19:Hα | |
| 50 | Val | 8.01 | 123.12 | 175.82 | 62.13 | - | | |
| 50 | Val | - | - | - | - | 32.37 | 3.75:Hα | |
| 51 | Lys | 8.18 | 125.70 | - | 56.22 | 32.61 | 3.95:Hα | |
| 51 | Lys | - | 125.62 | - | 56.30 | - | | |

| | | | | | | | | |
|---|---|---|---|---|---|---|---|---|
| 52 | Ser | 8.10 | 117.24 | 174.30 | 58.23 | 63.42 | 4.07:Hα | |
| 53 | Glu | 8.26 | 122.66 | 176.12 | 56.31 | 29.80 | 4.01:Hα | |
| 54 | Met | 8.06 | 120.19 | 175.88 | 55.04 | 32.67 | | |
| 54 | Met | - | - | - | - | - | 4.19:Hα | |
| 55 | Gly | 7.63 | 109.41 | - | 44.38 | - | 3.36:Hαa | |
| 56 | Pro | - | 133.45 | - | 63.07 | 31.39 | 3.95:Hα | |
| 57 | Trp | 7.65 | 119.90 | - | 57.40 | 28.48 | 4.25:Hα | |
| 58 | Met | 7.48 | 121.65 | 175.39 | 55.06 | 32.59 | 4.01:Hα | |
| 59 | Asp | 7.88 | 120.94 | 176.05 | 54.47 | 40.73 | 4.14:Hα | |
| 60 | Ser | - | - | 173.89 | - | - | | |
| 60 | Ser | 7.80 | 114.72 | - | 58.20 | 63.36 | 4.03:Hα | |
| 61 | Tyr | 7.89 | 122.18 | 175.49 | 58.01 | 38.48 | 4.22:Hα | |
| 62 | Ser | 7.92 | 118.16 | 173.87 | 57.65 | 63.70 | 4.15:Hα | |
| 63 | Gly | 7.10 | 109.79 | 177.60 | 44.50 | - | 3.64:Hαa | 3.64:Hαb |
| 64 | Pro | - | 133.23 | 176.85 | 63.14 | 31.47 | 3.99:Hα | |
| 65 | Tyr | 8.04 | 119.37 | 176.16 | 57.45 | 38.06 | 4.25:Hα | |
| 66 | Gly | 7.79 | 109.62 | 173.47 | 45.16 | - | 3.48:Hαa | 3.55:Hαb |
| 67 | Asp | 7.92 | 120.09 | - | - | - | | |
| 67 | Asp | - | - | 176.30 | 53.91 | 40.66 | 4.23:Hα | |
| 68 | Met | 8.08 | 121.07 | - | - | - | | |
| 68 | Met | - | - | 176.20 | 55.52 | 32.09 | | |
| 69 | Arg | 8.01 | 121.29 | 176.20 | 56.22 | 30.11 | 3.90:Hα | |
| 70 | Leu | 7.94 | 122.34 | - | - | - | | |
| 70 | Leu | - | - | 177.39 | 54.95 | 41.93 | 3.97:Hα | |
| 71 | Glu | 8.19 | 121.33 | 176.53 | 56.66 | 29.79 | 3.92:Hα | |
| 72 | Thr | 7.80 | 114.23 | 174.25 | 61.73 | 69.54 | 3.94:Hα | |
| 73 | Ala | 7.98 | 125.94 | 177.63 | 52.52 | 18.71 | 3.94:Hα | |
| 74 | Arg | 7.97 | 119.77 | 175.86 | 55.92 | 30.43 | 3.93:Hα | |
| 75 | Asp | - | - | 175.52 | 54.11 | 40.64 | 4.19:Hα | |
| 75 | Asp | 7.95 | 119.81 | - | - | - | | |
| 76 | His | 7.94 | 118.38 | 174.04 | - | 29.28 | | |
| 76 | His | - | - | - | 55.23 | - | 4.30:Hα | |
| 77 | Val | 7.85 | 121.58 | 175.62 | 62.01 | 32.33 | 3.71:Hα | |
| 78 | Leu | 8.12 | 127.56 | 174.73 | 52.51 | 41.22 | 4.22:Hα | |
| 79 | Pro | - | 135.18 | 176.82 | 62.61 | 31.72 | 4.07:Hα | |
| 80 | Ile | - | - | 175.57 | 61.51 | - | | |
| 80 | Ile | 7.96 | 120.28 | - | - | 38.44 | 3.73:Hα | |
| 81 | Asp | 7.96 | 121.73 | 175.47 | 53.59 | 40.55 | 4.16:Hα | |
| 82 | Tyr | 7.59 | 120.34 | 174.80 | 58.41 | 38.69 | 3.94:Hα | |
| 83 | Tyr | 7.63 | 121.13 | - | - | - | | |
| 83 | Tyr | - | - | 173.97 | 57.53 | 38.67 | 4.05:Hα | |
| 84 | Phe | 7.59 | 123.87 | 178.16 | 54.68 | 38.89 | 4.38:Hα | |
| 85 | Pro | - | 137.62 | - | 60.88 | - | 4.21:Hα | |
| 86 | Pro | - | 134.99 | 176.64 | 62.54 | 31.72 | 4.15:Hα | |
| 87 | Gln | 8.27 | 120.86 | 175.61 | 55.25 | 29.39 | 3.95:Hα | |
| 88 | Lys | 8.23 | 123.91 | 175.66 | 56.14 | 32.76 | 4.05:Hα | |
| 89 | Thr | 7.64 | 121.31 | 178.99 | 62.95 | 70.36 | 3.82:Hα | |
